# Cellular rules underlying psychedelic control of prefrontal pyramidal neurons

**DOI:** 10.1101/2023.10.20.563334

**Authors:** Tyler G Ekins, Isla Brooks, Sameer Kailasa, Chloe Rybicki-Kler, Izabela Jedrasiak-Cape, Ethan Donoho, George A. Mashour, Jason Rech, Omar J Ahmed

**Affiliations:** Dept. of Psychology, University of Michigan, Ann Arbor, MI 48109; Dept. of Mathematics, University of Michigan, Ann Arbor, MI 48109; Neuroscience Graduate Program, University of Michigan, Ann Arbor, MI 48109; Michigan Psychedelic Center, University of Michigan, Ann Arbor, MI 48109; Department of Medicinal Chemistry, University of Michigan, Ann Arbor, MI 48109; Dept. of Biomedical Engineering, University of Michigan, Ann Arbor, MI 48109

## Abstract

Classical psychedelic drugs are thought to increase excitability of pyramidal cells in prefrontal cortex via activation of serotonin 2_A_ receptors (5-HT2_A_Rs). Here, we instead find that multiple classes of psychedelics dose-dependently suppress intrinsic excitability of pyramidal neurons, and that extracellular delivery of psychedelics decreases excitability significantly more than intracellular delivery. A previously unknown mechanism underlies this psychedelic drug action: enhancement of ubiquitously expressed potassium “M-current” channels that is independent of 5-HT2R activation. Using machine-learning-based data assimilation models, we show that M-current activation interacts with previously described mechanisms to dramatically reduce intrinsic excitability and shorten working memory timespan. Thus, psychedelic drugs suppress intrinsic excitability by modulating ion channels that are expressed throughout the brain, potentially triggering homeostatic adjustments that can contribute to widespread therapeutic benefits.

## INTRODUCTION

Classical, or serotonergic, psychedelic drugs induce radical changes in perception and cognition, including altered working memory capacity (*1–10*). Psychedelics are being used successfully to treat neuropsychiatric disorders (*11–18*), but the underlying mechanisms remain unclear. Leading models posit that classical psychedelics acutely activate serotonin 2_A_ receptors (5-HT2_A_Rs) to increase excitability (Fig. 1A) of prefrontal cortex (PFC) pyramidal neurons (*19*–*24*), eventually resulting in a lasting enhancement of synaptic density (*25–31*). Neuronal excitability consists of two primary components: intrinsic electrical properties and synaptic input (*32–36*), with psychedelic enhancement of intrinsic excitability attributed to resting membrane potential (RMP) depolarization and reduction of spike afterhyperpolarization amplitude (*37–42*). However, these findings do not capture all the effects that serotonergic psychedelics have on cellular neurophysiology; it has also been reported that psychedelics escalate inactivation of transient sodium channels (*43*, *44*). Thus, it is unknown if serotonergic psychedelics universally enhance intrinsic excitability of PFC pyramidal cells (PC).

**Fig. 1.**
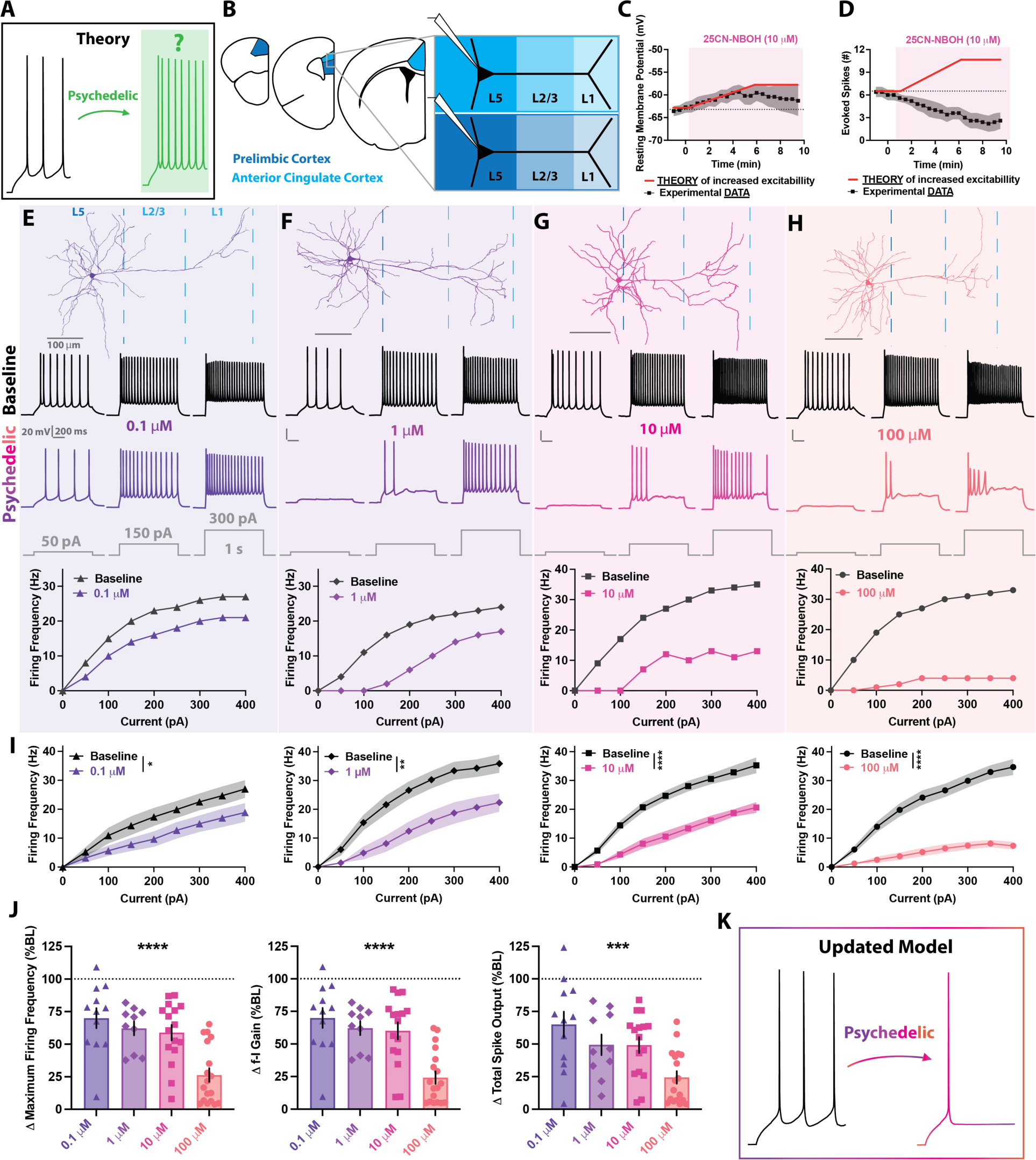
Psychedelic-induced suppression of intrinsic excitability. (A) Leading current theories posit that psychedelics increase spiking output of L5 PCs. (B) PFC PCs were recorded in L5 of Prelimbic and Anterior Cingulate Cortex. (C) Consistent with part of the theory of increased excitability, the psychedelic drug 25CN-NBOH (NBOH) depolarized resting membrane potentials by a few millivolts. (D) Contrary to current theory, NBOH suppressed evoked spiking. (E to H) Individual PFC L5 PC morphological reconstructions (top) and corresponding firing patterns (middle) and f-I curves (bottom) during baseline (black) and after 10-minute bath perfusion of NBOH (color): 0.1 µM (E), 1 µM (F), 10 µM (G), 100 µM (H). (I) Mean f-I curves at baseline and in NBOH, showing that NBOH significantly suppresses spiking, with more suppression at higher doses. (J) NBOH strongly attenuated maximum firing frequency, f-I gain and total spike output, and this suppression of spiking was strongly dose-dependent. (K) The revised framework for understanding psychedelic action indicates that psychedelics suppress rather than enhance overall intrinsic excitability and spiking output of PFC L5 PCs. *p<0.05; **p<0.01; ***p<0.001; ****p<0.0001, F-test between simple linear regressions of baseline and drug (I), one-way ANOVA of %BL changes between dose groups (J). Error bars and shaded regions represent standard error of the mean (SEM).

Here, we use whole-cell electrophysiology, morphology, pharmacology, machine-learning based data assimilation, and computational modeling to test the hypothesis that serotonergic psychedelics increase both intrinsic and synaptic excitability of PFC PCs through activation of extracellular 5-HT2_A_Rs.

## RESULTS

### Psychedelics dose-dependently suppress intrinsic excitability of PFC Layer 5 PCs

We first investigated how the serotonergic psychedelic and 5-HT2_A_R-preferring agonist 25CN-NBOH (*45–50*) alters RMPs and evoked firing rates of PFC layer 5 (L5) PCs (Fig. 1B). This experiment utilized no holding current, allowing cells to sit at rest between depolarizing step pulses. Consistent with previous studies (*38–41*), we found psychedelic-mediated RMP depolarization (Fig. 1C). However, this depolarization was accompanied by a seemingly paradoxical decrease in the number of evoked spikes, suggestive of an overall suppression of intrinsic excitability (Fig. 1D).

To better understand this ostensible paradox, we applied NBOH for 10 minutes at four dose levels: 0.1, 1, 10 and 100 µM, and assessed intrinsic membrane and firing properties both pre- and post-exposure. Again, contrary to predictions based on current theories, NBOH robustly suppressed evoked spiking of PFC L5 PCs (Fig. 1E-H). All doses significantly decreased maximum firing frequency, frequency-current (f-I) gain and total spiking output, and this suppression of intrinsic excitability was strongly dose-dependent (Fig. 1I-J; table S1). Importantly, this psychedelic-induced spiking suppression was observed across the following parameters: anteroposterior location within PFC, somatic position within L5, age, genotype, and sex (fig. S1).

Unlike the intrinsic excitability changes, alterations in synaptic excitability were consistent with the current consensus, albeit with important caveats related to dose. We found that NBOH dose-dependently enhanced sEPSC frequency in PFC L5 PCs (fig. S2A-E). A sustained frequency elevation was observed only at higher doses (10 and 100 µM), indicating a thresholded, dose-dependent enhancement of spontaneous glutamate release (fig. S2F; table S2). To determine if these intrinsic and synaptic excitability changes generalize to other classes of serotonergic psychedelic drugs, we repeated these experiments with the psychedelic phenethylamine DOI and psychedelic tryptamine 4-HO-DiPT (fig. S3). Similar to NBOH, both DOI and 4-HO-DiPT dose-dependently suppressed intrinsic excitability (fig. S3A-F) and enhanced synaptic excitability (fig. S3G). Thus, serotonergic psychedelics of different classes induce similar dose-dependent changes on intrinsic and synaptic neurophysiology of PFC L5 PCs (Fig. 1K; table S1-2).

It has been proposed that intracellular 5-HT2_A_Rs may contribute to the subjective effects of psychedelic drugs (*27*). Intracellular 5-HT2_A_Rs are, however, not responsible for the enhancement of synaptic excitability, which also occurs upon activation by the membrane-impermeable agonist serotonin and is dependent on blockade of presynaptic axonal Kv1.2 channels (*39*, *51–60*). If intracellular 5-HT2_A_Rs, activated by psychedelic drugs diffusing into neuronal membranes, are responsible for the changes to intrinsic excitability, then intracellular delivery of psychedelics should suppress intrinsic excitability to the same extent as an equivalent bath-applied concentration (fig. S4A). To test this hypothesis, we replicated our earlier experiments, intracellularly administering 1, 10 or 100 µM NBOH directly through the patch electrode (fig. S4B-E).

Intracellular delivery of psychedelics was significantly less effective at suppressing intrinsic excitability compared to extracellular delivery (fig. S4E,G). As NBOH is brain penetrant and membrane permeable (*45*), some percentage of the drug likely diffuses out of the cell membrane where it can activate extracellular 5-HT2Rs located on somatodendritic compartments of the recorded neurons and on presynaptic terminals of input axons. In support of extracellular 5-HT2_A_R activation, we observed a sustained sEPSC frequency enhancement in 33% of cells with intracellular delivery of 100 µM NBOH, and only 15% of cells with intracellular delivery of 10 µM (fig. S4F). However, as was the case with intrinsic excitability, this effect was significantly weaker than with extracellular delivery (62% with 100 µM, 55 % with 10 µM; fig. S4F-G). Therefore, serotonergic psychedelics dose-dependently suppress intrinsic excitability and enhance synaptic excitability of PFC L5 PCs with minimal indication of the involvement of intracellular 5-HT2_A_Rs (fig. S4H; table S1-2).

### Excitability suppression mechanisms are dose-dependent

We next explored if the mechanisms of spike loss were also dose-dependent (Fig. 2A). At low to moderate doses, we noticed that post-rheobase spike loss was often due to increased spike frequency adaptation (Fig. 2B). In contrast, the highest dose often resulted in further spike loss due to depolarization block (Fig. 2C).

**Fig. 2.**
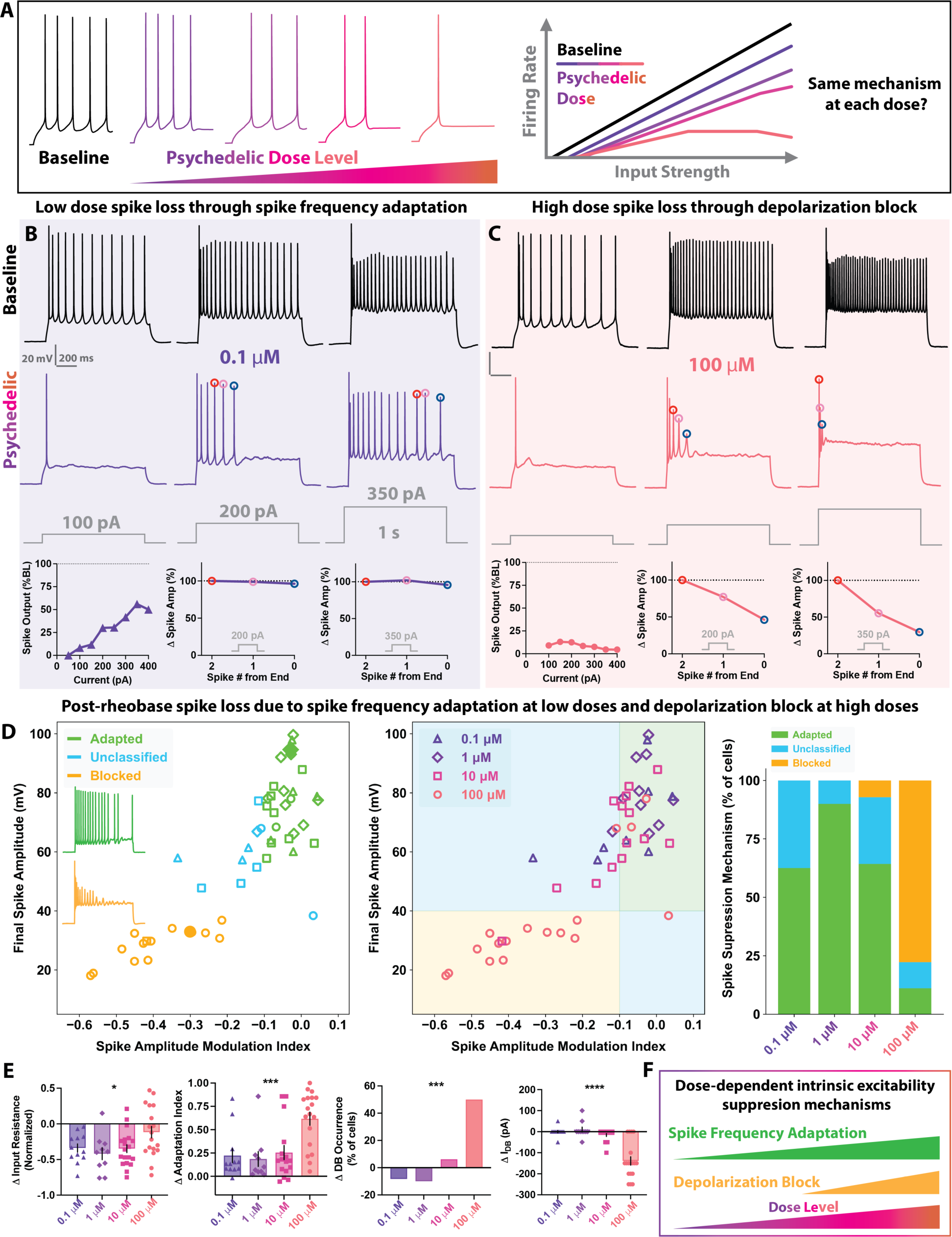
Dose-dependent mechanisms for intrinsic excitability suppression. (A) Model of dose-dependent suppression of spiking by psychedelic drugs. Higher doses cause stronger suppression, but is the mechanism the same? (B-C) Dose-dependent spike loss through spike frequency adaptation (B) or depolarization block (C). Individual PFC L5 PC firing patterns (top) during baseline (black) and after 10-minute bath perfusion of NBOH (color). f-I curves showing spiking output as % of baseline spiking (bottom left). Spike amplitude of the last 3 spikes plotted as % of the 3^rd^ to last spike for the current step shown above: 200 pA (bottom middle), 350 pA (bottom right). Note that the spike amplitudes remained constant for the 0.1 µM cell that exhibited increased spike frequency adaptation and that the spike amplitudes rapidly decreased for 100 µM cell that entered depolarization block. (D) Spike amplitude modulation plots identify signatures of spike loss due to spike frequency adaptation or depolarization block (left). A filled green diamond corresponds to the top (green) inset firing trace and bottom filled circle corresponds to the bottom (orange) inset firing trace, cells that are respectively adapted or blocked by NBOH. Spike amplitude modulation plot overlaying effect (background color) and dose (symbol shape/color), and dose summary plots indicate dose-dependent spike loss through spike frequency adaptation or depolarization block (middle, right). (E) Psychedelics induce dose-dependent changes to input resistance, spike frequency adaptation index, depolarization block occurrence, and current to induce depolarization block. (F) Psychedelic drugs decrease intrinsic excitability through enhancing spike frequency adaptation at low to moderate doses and increasing depolarization block susceptibility at high doses. *p<0.05; **p<0.01; ***p<0.001; ****p<0.0001, one-way ANOVA of %BL changes between dose groups (E). Error bars represent SEM.

To quantitatively determine the dominant spike loss mechanism for each psychedelic-suppressed cell, we analyzed the final spike amplitude and the modulation index of spike amplitudes (see methods). Cells with large final spike amplitudes and modulation indices close to 0 were classified as adapted, whereas cells with small final spike amplitudes and negative modulation indices were classified as in depolarization block (blocked). Unclassified cells had final amplitudes and modulation indices between the two other groups (Fig. 2D). We confirmed our initial observation: low to moderate doses primarily enhanced spike frequency adaptation, whereas the highest dose increased the likelihood of depolarization block (Fig. 2D-E). Interestingly, low to moderate doses lowered input resistance more than the highest dose, potentially indicating dose-dependent conductance changes (Fig. 2E).

We repeated this analysis for DOI and 4-HO-DiPT and found similar results (fig. S5A). Given the consistent dose-dependent impacts on intrinsic excitability irrespective of the specific serotonergic psychedelic used, we pooled datasets to examine correlations of excitability suppression with other factors (fig. S5B). Dose-dependent mechanism-specific spike loss was again observed throughout sexes, positions within PFC, laminar depths within L5, or ages (fig. S5C-F). Hence, serotonergic psychedelics suppress intrinsic excitability of PFC L5 PCs through distinct cellular mechanisms: dose-dependent spike frequency adaptation that is seen even at the lowest doses and thresholded dose-dependent depolarization block that emerges at higher doses (Fig. 2F).

### Psychedelics decrease excitability by enhancing potassium M-current independently of 5-HT2R activation

Depolarization block involves the progressive inactivation of sodium channels and can be explained by previously identified 5-HT2R-mediated changes to transient sodium conductance (*43*, *44*). However, sodium channel inactivation does not explain the input resistance decreases, rheobase increases, and other changes to intrinsic excitability elicited by psychedelic drugs (table S1). To uncover the additional ion channel mechanisms of psychedelic-induced hypoexcitability that are seen even at the lowest psychedelic doses, we utilized pharmacology-assisted whole-cell electrophysiology. Changes to firing patterns, in particular resulting from low to moderate doses of psychedelics, resembled firing changes in response to potassium-channel-mediated M-current activation (*61*). Therefore, we hypothesized that psychedelics might activate 5-HT2Rs to initiate a signaling cascade to enhance activity of M-current, which suppresses intrinsic excitability (Fig. 3A).

**Fig. 3.**
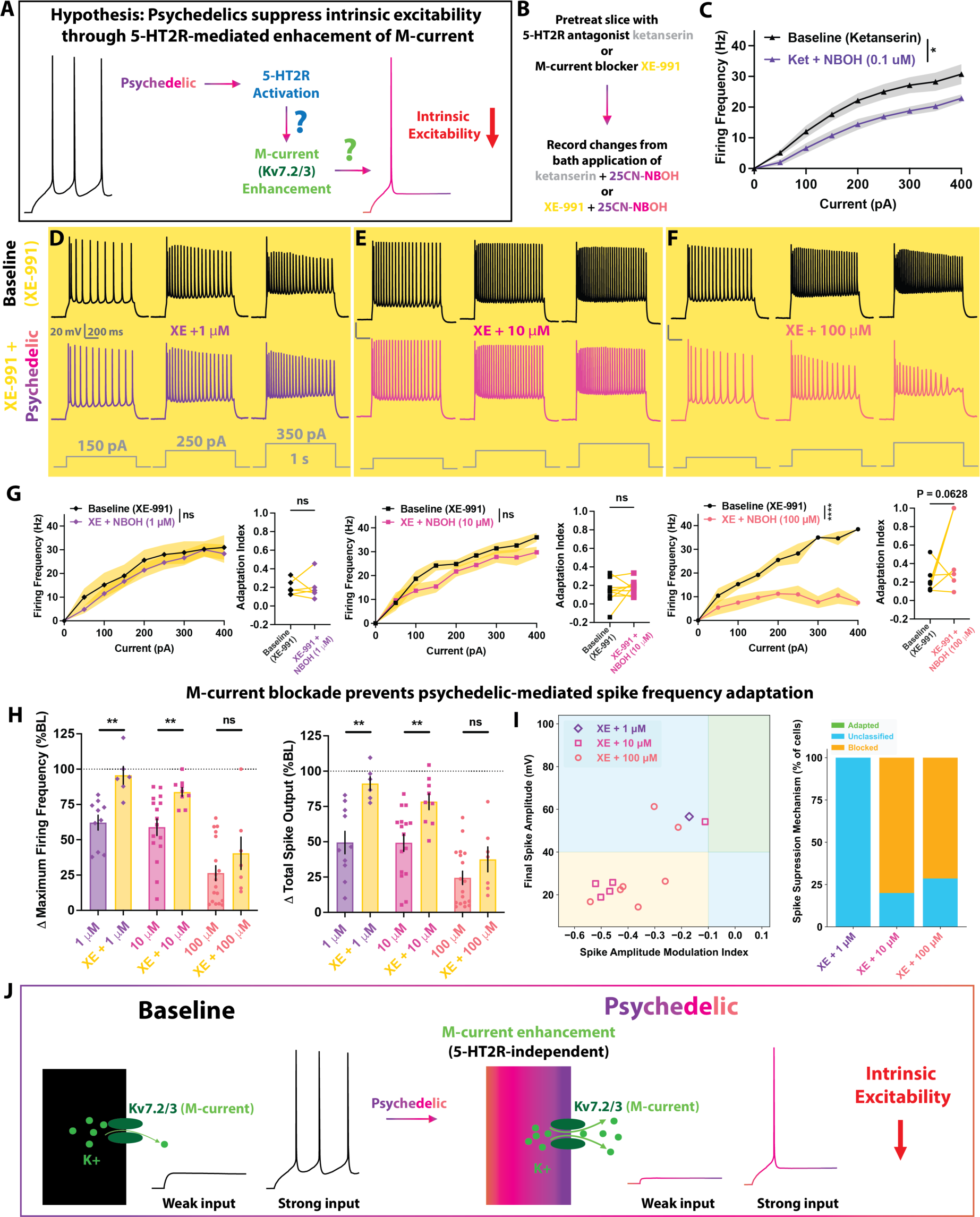
Psychedelics suppress excitability through 5-HT2R-independent M-current activation. (A) 5-HT2Rs are considered to be the primary target of psychedelic drugs. We tested the hypothesis that psychedelics may suppress intrinsic excitability and increase spike-frequency adaptation through 5-HT2R-mediated enhancement of M-current/Kv7). (B) Experimental design. Slices are pretreated with 5-HT2R antagonist ketanserin (10 µM) or the M/Kv7 blocker XE-991 (10 µM), after which baseline recordings are taken. Changes to evoked spiking are measured following a 10-minute perfusion of the antagonist/blocker in combination with varying doses of NBOH. (C) Ketanserin pretreatment does not prevent 0.1 µM NBOH-induced suppression of spiking. (D to F) Individual PFC L5 PC firing patterns in the presence of XE-991 (yellow background) during baseline (black) and after 10-minute bath perfusion of NBOH (color): 1 µM (D), 10 µM (E), 100 µM (F). (G) XE-991 pretreatment prevents 1 and 10, but not 100 µM NBOH-induced suppression of spiking and increase of spike frequency adaptation. (H) Blocking M/Kv7 strongly attenuates the ability of NBOH to suppress spiking at doses that normally promote adaptation-induced spike loss (1 and 10 µM). (I) Spike amplitude modulation plots and summary plots for the cells exhibiting spike suppression under XE-991 + NBOH, indicating that spike loss is not due to adaptation. (J) Model of psychedelic induced 5-HT2R-independent suppression of intrinsic excitability via M-current enhancement. NS, not significant; *p<0.05; **p<0.01; ***p<0.001; ****p<0.0001, F-test between simple linear regressions of baseline (XE-991) and drug (XE-991 + NBOH) f-I curves or two-tailed paired t test between baseline and drug adaptation index (G), two-tailed unpaired t test or Mann Whitney test between baseline and drug (H). Error bars represent SEM.

To test the hypothesis that intrinsic excitability suppression is dependent on 5-HT2Rs, we blocked 5-HT2Rs with bath application of ketanserin (10 µM) before introducing NBOH along with ketanserin (Fig. 3B). Notably, even with ketanserin pretreatment, the psychedelic-induced hypoexcitability was not eliminated (Fig. 3C; fig. S6A; table S3). This suggests that psychedelics can reduce intrinsic excitability without the requirement of 5-HT2Rs. Ketanserin pretreatment did, however, prevent the NBOH-mediated enhancement of sEPSCs (fig. S6B; table S4), consistent with the known involvement of 5-HT2Rs in increasing synaptic event frequency (*39*, *51*, *52*, *55*, *62*, *63*).

We next investigated whether psychedelics enhance M-current activation. To examine this, we blocked M-current/Kv7 channels with XE-991 (10 µM) and subsequently introduced NBOH along with XE-991 (Fig. 3B). Unlike ketanserin, XE-991 pretreatment eliminated psychedelic-induced enhancement of spike frequency adaptation and significantly attenuated suppression of firing at low to moderate doses (Fig. 3D-H; table S3). Again, in contrast to ketanserin, XE-991 pretreatment did not block NBOH-mediated sEPSC frequency enhancement (fig. S6C; table S4). We found that of the neurons that still underwent psychedelic spike suppression, none were classified as adapted (Fig. 3I). Therefore, serotonergic psychedelics are able to suppress intrinsic excitability through 5-HT2R-independent enhancement of M/Kv7 conductance (Fig. 3J).

### M-current and transient sodium current alterations combine to suppress spiking and impair cellular correlates of persistent working memory

In order to further verify that the proposed channel mechanisms – upregulation of M/Kv7 conductance concomitant with known effects on transient sodium channels (*43*, *44*) – could account for the experimentally observed psychedelic-induced hypoexcitability, we sought to construct a computational model to reproduce the experimental observations (Fig. 4A). Using a recently developed machine-learning-based data assimilation framework (*64*, *65*), we built a conductance-based Hodgkin-Huxley model capable of predicting the voltage response to current stimuli unseen by the assimilation process (Fig 4B). Thus, we may consider the model to be a reasonable approximation for the input-output transform implemented by the actual cortical neuron.

**Fig. 4.**
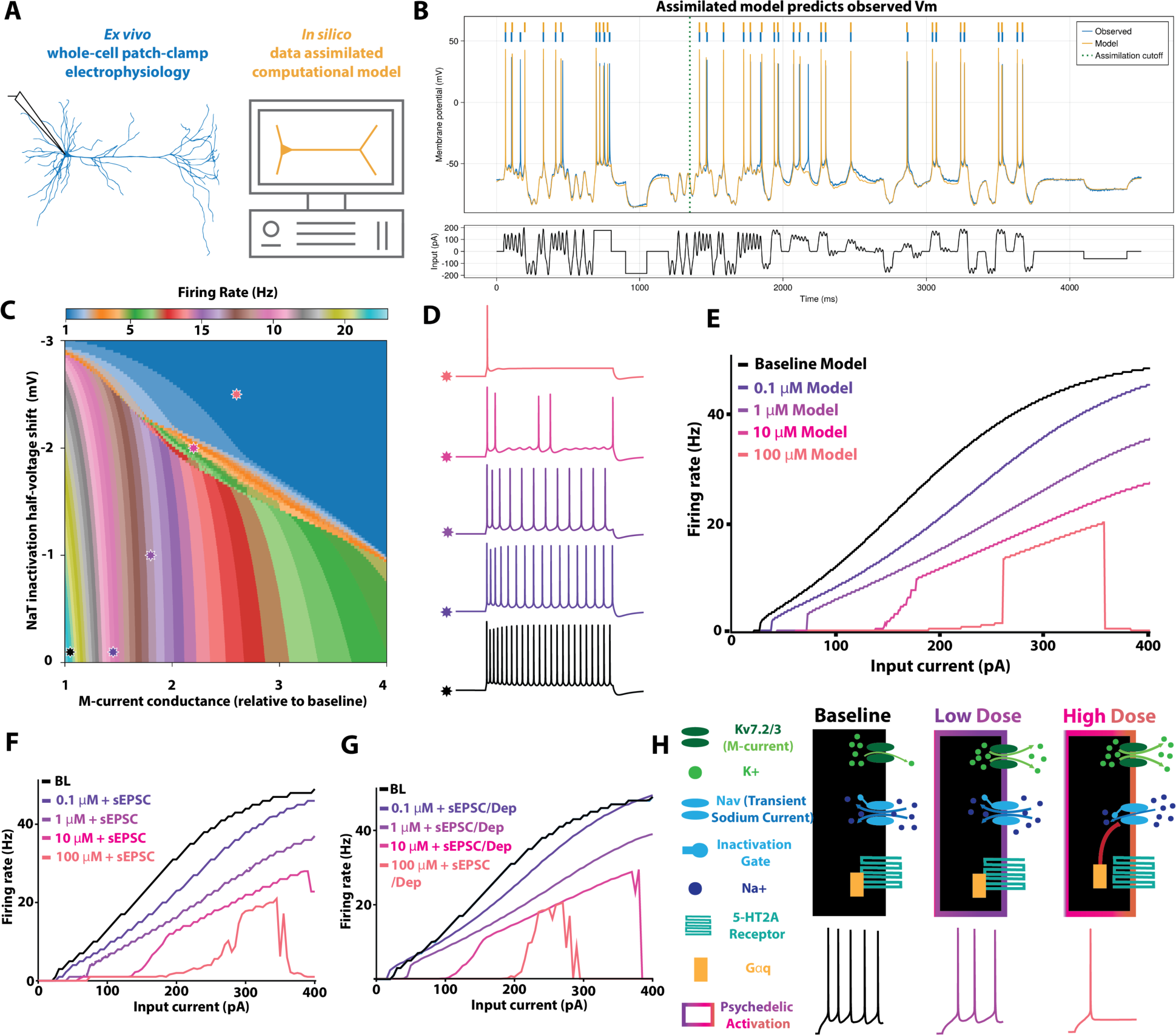
Machine-learning-based data assimilation model of psychedelic drug action reveals spike suppression. (A) Experimental overview. A predictive PFC L5 PC model was assimilated with machine learning and current clamp data. (B). The assimilated model captures dynamical features of the neuron. Voltage traces of the real (blue) and assimilated (orange) neurons are displayed. Spike times are overlaid (top) and the chaotic input current used for the assimilation is shown (bottom). The cutoff of the assimilation is shown as a dotted line. (C) Increasing M/Kv7 conductance and hyperpolarizing the inactivation half-voltage of transient sodium current (NaT) decrease evoked spiking. Manipulating both simultaneously enhances spiking suppression. Plot indicates firing rates for a 1-second, 160 pA current step; stars, from left to right, indicate the baseline and proposed psychedelic dose models. (D) Firing traces of the assimilated neuron at baseline and the 4 proposed psychedelic dose models (Baseline: M/Kv7 conductance (gM) =0.05, change in NaT inactivation half-voltage (ΔθH) =0.0; 0.1 µM: gM=0.07, ΔθH=0.0; 1 µM: gM=0.09, ΔθH=-1.0; 10 µM: gM=0.11, ΔθH=-2.0; 100 µM: gM=0.13, ΔθH=-2.5). (E) f-I curves of the assimilated neuron at baseline and the 4 psychedelic dose models. (F) f-I curves of the assimilated neuron with added simulated sEPSCs mimicking the effects observed at each dose. (G) f-I curves of the assimilated neuron with added sEPSCs and RMP depolarization mimicking the effects observed at each dose. (H) Ion channel model of psychedelic action in PFC L5 PCs. At low doses, psychedelics enhance M-current through a 5-HT2R-independent mechanism. This results in decreased excitability through enhancement of spike frequency adaptation (middle). At higher doses, psychedelics additionally recruit 5-HT2Rs to initiate a signaling cascade (represented by red curve) to modulate transient sodium currents. The combination of enhancing M-current and hyperpolarizing the inactivation half-voltage of transient sodium current results in radical spike suppression at higher doses.

Proceeding with this model, we performed *in silico* perturbation experiments, varying the maximal conductance of M/Kv7 current and the inactivation half-voltage of transient sodium current (NaT). As expected, increases in the maximal conductance of M/Kv7, or hyperpolarization of the half-inactivation voltage of NaT lead to a graded decrease in spike output (Fig. 4C-E). We observed similar results when additionally varying the maximal conductance of NaT (data not shown). Intriguingly, concomitant manipulations of both M/Kv7 and NaT to certain levels can transition the assimilated model into an irregular firing mode (*66*). In this regime, spiking output alternates with sustained subthreshold oscillations, despite the application of a constant input current. Within our own experimental dataset, we observe putative examples of such firing patterns induced by moderate doses of psychedelics (fig. S8). Once again, in alignment with experimental data, we observed depolarization block in the 100 µM model (Fig. 4E; fig. S7).

Next, we sought to examine which of the two experimentally observed effects would dominate in the computational model: the increase in synaptic excitability, via increased sEPSC frequency, or the decrease in intrinsic excitability via proposed channel mechanisms. We simulated the model cell under conditions of bombardment by stochastic sEPSCs, with kinetics, amplitude, and frequency matched to our experimental data (Table 5). We found that presence of stochastic sEPSCs minimally modified the f-I relation of the model cell, indicating that the change in intrinsic excitability dominates the model’s input-output relation (Fig. 4F). We repeated these experiments by additionally incorporating the experimentally observed RMP depolarization (Table 5; Fig. 4G) and again observed analogous results: dose-dependent spike suppression that was seen even at the lowest psychedelic dose models. Finally, to confirm that the observed effects were not specific to our machine-learning-based assimilated model, we repeated *in silico* experiments with a minimal neuron model that has been frequently used in computational studies (*67*) and found similar results (fig. S9).

Lastly, as psychedelics have been shown to dose-dependently reduce working memory capacity (*7–10*), we modeled the impact of psychedelic modulation of intrinsic excitability on known cellular correlates of working memory. To achieve this, we first added a long-lasting calcium-activated nonspecific cation conductance (I_CAN_) to our assimilated model neuron (fig. S10). Upon adding this additional conductance, stimulus-induced spiking results in continuous spiking that outlives the stimulus (fig. S10A,B), a cholinergic activation-dependent cellular correlate of working memory known as ‘persistent firing’ (*68–71*). Psychedelic modulation dose-dependently disrupted persistent firing, reducing the number of persistent spikes (0.1 µM), causing self-termination of spiking (1 µM), or eliminating the ability to produce any persistent spikes (10-100 µM; fig. S10A-D).

Overall, our computational results are consistent with the hypothesis that concomitant and graded modulation of M/Kv7 and NaT current result in dose-dependent intrinsic excitability suppression (Fig. 4H). Moreover, within our experimentally constrained modeling framework, these reductions in intrinsic cell excitability through active conductance modulations override the observed increases in synaptic excitability and resting membrane potential depolarization, resulting in a net blunting of cell output, providing a mechanistic complement to similar experimental observations of spike suppression (Fig. 1D).

## DISCUSSION

Although psychedelics do enhance some forms of neuronal excitability, these data indicate that the dominant and net effect is dose-dependent suppression of intrinsic excitability, necessitating a revision of the current paradigm (*19–24*). Our acute results differ from some previous studies that concluded that psychedelics increase intrinsic excitability, likely due to prior studies examining RMP changes or using only near-threshold current injections, which fail to reveal the larger intrinsic suppression changes (*37*, *38*, *40*, *42*). Additionally, we utilized psychedelic drugs (*47–50*, *72–76*), in contrast to many former studies that used 5-HT2AR agonists with non- or unknown psychedelic effects such as serotonin or alpha-methyl-serotonin (*77–84*).

These results establish a set of fundamental cellular rules for psychedelic control of neuronal activity. 5-HT2R-independent activation of M/Kv7 conductance suggests that serotonergic psychedelics suppress intrinsic excitability throughout the entire brain in neurons with prominent Kv7 channels, such as neocortical and hippocampal PCs (*61*, *85*). Higher doses, which strongly activate 5-HT2_A_Rs in addition to M/Kv7, will further suppress spiking of PFC L5 PCs, due to dense expression of 5-HT2_A_Rs in these cells (*85–88*). Although firing rates *in vivo* are controlled by numerous variables, making it difficult to ascertain the direct influence of decreased intrinsic excitability, supporting evidence is found in several *in vivo* electrophysiology studies in which serotonergic psychedelics have mixed effects on firing rates but result in a net firing decrease throughout the brain, including dose-dependent suppression in PFC (*89–100*).

Acute spiking suppression with enhanced spontaneous glutamate release may contribute to the lasting synaptic changes to PFC L5 PCs elicited by single *in vivo* doses of psychedelics observed by others (*25–27*) and us (fig. S11), through post-silencing rebound synaptogenesis in combination with intracellular 5-HT2_A_R-mediated spinogenesis (*27*, *101–104*), possibly coinciding with a temporary reversion to a potentially therapeutic critical period-like state (*105*– *107*). Thus, dose-dependent spiking suppression of PFC PCs may contribute to subjective psychedelic effects and lasting therapeutic outcome (fig. S12).

## Author contributions

Conceptualization: TGE, IB, SK, OJA

Methodology: TGE, IB, SK, CRK, IJC, ED, JR, OJA

Investigation: TGE, IB, SK, CRK, IJC, ED

Visualization: TGE, IB, SK, OJA

Project administration: OJA Supervision: OJA

Writing – original draft: TGE, IB, SK, OJA

Writing – review & editing: TGE, IB, SK, CRK, IJC, ED, GM, JR, OJA

## SUPPLEMENTAL INFORMATION

### MATERIALS AND METHODS

#### Mice

All procedures and use of animal were approved by the University of Michigan Institutional Animal Care and Use Committee. The following mouse lines were used for experiments: C57BL/6J (Jackson Laboratories #000664), Chat-IRES-Cre (Jackson Laboratories #006410), Cux2-Cre (MMRRC #032778-MU), Grp-Cre-KI (#033174), Nex-Cre (*108*), B6 Pvalb-IRES-Cre (Jackson Laboratories #017320), Scnn1a-Tg3-Cre (Jackson Laboratories #009613), Syt6-Cre (MMRRC #032012-UCD), Trpv1-Cre (Jackson Laboratories #017769), Ai14 (Jackson Laboratories #007914), Kj319-Sla-Cre::Ai14 (In-house crossing of Ai14 (Jackson Labs) with Kj319-Sla-Cre, MMRRC #033066-UCD), Ai32 (Jackson Laboratories #024109), Nex-Cre::Ai32 (In-house crossing of Ai32 with Nex-Cre), Scnn1a-Tg3-Cre::Ai32 (In-house crossing of Scnn1a-Tg3-Cre with Ai32). A total of 73 mice (185 cells) of both sexes between postnatal days 22-241 were used in this study. 69 mice (162 cells) were used for acute drug recordings and 4 mice (23 cells) were used for intrinsic and synaptic characterization after *in vivo* drug injection.

#### Drugs

All drugs were dissolved in water before mixing into artificial cerebrospinal fluid for *ex vivo* experiments or sterile saline for *in vivo* experiments. DOI was purchased from Sigma-Aldrich. 4-HO-DiPT was purchased from Cayman Chemical. 25CN-NBOH, XE-991 and Ketanserin were purchased from Tocris Bioscience. Sonication was used to assist solubilization of XE-991 and Ketanserin in water. Ketanserin was chosen in place of more ‘selective’ 5-HT2_A_R antagonists such as MDL,100907 (*109*) due to solubility. DMSO was avoided as a solvent as it is known to alter intrinsic electrophysiological properties of neocortical and hippocampal neurons at doses far lower than regularly used in neuroscience experiments (*110*).

#### Slice Preparation

Mice underwent deep isoflurane anesthesia before decapitation. Brains were dissected out in ice-cold sucrose-substituted ACSF, saturated with 95% O_2_ and 5% CO_2_ and containing the following (in mM): 3 KCl, 1.25 NaH2PO4, 26 NaHCO3, 10 dextrose, 234 sucrose, 0.2 CaCl2, 4 MgSO4. 300 um coronal slices were cut using a 1200VT vibratome (Leica Microsystems) and placed in a high-magnesium ACSF solution at 32 ° C for 30 minutes, after which they rested at room temperature for at least 30 more minutes before being recorded. During experiments, slices were submerged in a recording chamber with physiological temperature (32 ° C) ACSF (126 mM NaCl, 1.25 mM NaH2PO4, 26 mM NaHCO3, 3 mM KCl, 10 mM dextrose, 1.20 mM CaCl2, and 1 mM MgSO4) perfused at a rate of 3 mL/minute. Slices containing prelimbic cortex and anterior cingulate cortex were obtained from anteroposterior distance to bregma 2.9 to -0.5 mm. Infralimbic cortex was not targeted for recording due to reports of lower Htr2a expression (*88*, *111–113*).

#### Layer 5 regular spiking pyramidal cell recording and quality control

Neurons were visualized on an Olympus BX51WI microscope, Olympus 60x water immersion lens, and Andor Neo sCMOS camera (Oxford Instruments, Abingdon, Oxfordshire, UK). Patch electrodes were pulled from borosilicate glass (Sutter Instruments) and had a diameter of 2-4 µm and resistances between 3-5 MΩ. The internal solution was potassium gluconate-based and contained (in mM): 130 K-gluconate, 0.6 EGTA, 10 HEPES, 2 MgATP, 0.3 Na2GTP, 6 KCl, NaCl and 0.5% biocytin (calculated chloride reversal potential of -68 mV; pH of 7.25; osmolarity of 290 mOsm). For intracellular drug delivery experiments, 25CN-NBOH (1, 10, or 100 µM) was additionally added to the internal solution. Pipette capacitance compensation and bridge balance were applied; recordings were not corrected post-hoc for liquid junction potential. Only neurons with regular spiking firing patterns were used in this study. Cells were additionally excluded if baseline resting membrane potential was more depolarized than -50 mV, if input resistance decreased > 25% before acute pharmacology experiments, if uncompensated series resistance (Rs) was > 35 MΩ, or if Δ Rs > 25% during acute pharmacology experiments. All whole-cell recordings were conducted with MultiClamp 700B amplifier and digitized at 20 kHz with Digidata 1550B (Molecular Devices) for collection on a computer equipped with pClamp 10.7 software (Molecular Devices). Patch electrodes additionally contained biocytin, which enabled post-hoc imaging (see methods below) to determine somatic position and confirm pyramidal morphology. We considered L5 neurons to have somatic positions from 125 to 400 microns from the L1/2 boundary. However, as anterior-most regions of PFC have a considerably wider L5, some included neurons from those PFC subregions had deeper cell bodies (up to 625 microns from L1/2 border).

#### Current-clamp recordings and intrinsic property analysis

In the first experiment, no holding current was applied, and spikes were evoked through depolarizing square waves (0.5s, 50-100 pA) injected from rest every 30 seconds. After establishing a stable baseline spiking response (5 to 8 spikes, ±1 spike/cell) and RMP (-66 to -60 mV, ±2 mV/cell), 25CN-NBOH was bath-applied for 10 minutes and RMP and evoked firing rates continued to be recorded under identical stimulation parameters. For subsequent experiments, membrane potentials were biased to -65 mV at the start of each sweep. Firing patterns were investigated using a series of 1-second current injections (step size 50 pA) until 400 pA was injected or depolarization block was induced (with an exception for in-drug recordings in which the final current level was matched to the baseline maximal current). Firing frequency was the number of spikes that occurred for each 1-second current step. Spikes that did not have a peak voltage reaching at least -10 mV were not counted. F-I gain was the slope of a linear regression on the number of spikes at each current injection from 0 pA to the injection at which maximum firing frequency was attained, or 400pA, whichever came first. Total spike output was the sum of all spikes until maximum firing frequency or 400 pA. We used Python (version 3.10.4) to analyze all intrinsic properties, loading all ABF recording files with pyABF (*114*) and using the packages NumPy (*115*), pandas (*116*), and Matplotlib (*117*). For spike properties, we defined threshold as the voltage where the slope trajectory (dV/dT) reached 10 mV/ms. Amplitude was determined by the voltage difference between the threshold and the peak. Half-width was measured at the voltage corresponding to half of the spike amplitude. Maximum rise slope was the maximum of dV/dt; maximum decay slope was the minimum of dV/dt. These properties were measured for the first spike evoked by the 1-second current injection. Rheobase and latency to first spike were determined with a series of 2-second-long depolarizing current steps sufficient to induce spiking (5-10 pA step size), repeated 1-3 times. Latency to first spike was determined by measuring the interval between the onset of the current step and the time of the initial spike threshold voltage; rheobase was defined as the minimal current necessary to induce a spike during the 2-second current step. Adaptation index was computed at approximately twice the rheobase current from the 1-second current injection and was the number of spikes in the second half of the sweep subtracted from the number of spikes in the first half, divided by the total number of spikes. Additionally, we assessed input resistance (Rin), input capacitance (Cin), and the membrane time constant (TC) with a series of small hyperpolarizing current steps (-10 to -20 pA). Rin was computed using the voltage difference between the maximum response during the current injection and the mean voltage of the 100ms prior to the current injection, divided by the injected current. TC was computed by fitting an exponential decay curve to the voltage trace from the start of the current injection to the maximum voltage response. Input (membrane) capacitance (Cin) was calculated using the relationship Cin = TC/Rin. Injection of hyperpolarizing current into some neurons can produce a noticeable “sag”, which indicates presence of a hyperpolarization-activated cation current (I_h_) that activates after the initial hyperpolarization peak. Sag ratio was computed as the maximum response amplitude divided by the mean response amplitude of the last 50ms of the current injection. Response amplitudes were computed relative to the baseline voltage, computed as the mean of the 100ms prior to the current injection (*118–122*)

#### Classification of Spike Suppression Mechanisms

We classified individual neurons’ proposed mechanisms of psychedelic-induced spike loss based on their responses to 1-second current injections at 50pA steps between 50pA and a maximum of 400pA, taken both at baseline and in the drug condition. For each current injection in this range, the properties and total numbers of action potentials in the equivalent baseline and drug sweeps were analyzed. The highest three current injection values that were applied in both conditions were selected for the following analyses (e.g. if 50-350pA were applied at baseline, but 50-400pA in the drug condition, then the 250, 300, and 350pA sweeps were selected for both conditions).

For each selected sweep, putative action potentials were detected using Scipy’s find_peaks function, with a minimum prominence of 10mV and minimum distance of 2ms. Next, the prominence of each action potential, defined as the distance between the peak and the more positive of the preceding and following minima in the voltage trace, was computed. The maximum slope of each action potential, defined as the highest value of the derivative of voltage with respect to time during the action potential, was additionally computed. Only action potentials with a maximum slope of at least 10mV/ms, a distance between left and right troughs of less than 100ms, and a half-prominence width of less than 60ms were included in downstream analysis.

For each baseline-drug pair of selected current injections, the spike loss modulation index was computed as the number of spikes at baseline, minus the number of spikes in the drug condition, divided by the total number of spikes across both conditions. For each selected sweep in the drug condition, the spike amplitude modulation index was computed as the prominence of the final spike in the sweep, minus the prominence of the spike at the 33^rd^ percentile of sequentially assigned spike indices, divided by the prominence of the same 33^rd^ percentile spike.

Only sweeps with a spike loss modulation index of at least 0.1 were included in the following analysis. Cells with a mean spike amplitude modulation index of less than -0.1 and a mean final spike amplitude of less than 40mV across selected sweeps were classified as Blocked, whereas cells with a mean spike amplitude modulation index of at least -0.1 and a mean final spike amplitude of at least 40mV were classified as Adapted. Cells that met neither of these two criteria were marked Unclassified.

#### Voltage-clamp recordings and sEPSC analysis

All voltage-clamp recordings were conducted at a holding potential of -70 mV. sEPSC recordings were taken in 30-second sweeps with a brief (250 ms) hyperpolarizing test pulse (-5 mV) at the start to monitor Rs and Rin throughout the experiment. For standard pharmacology experiments, sEPSCs are recorded for two consecutive sweeps in standard ACSF, immediately before switching to the drug-containing ACSF and recording sEPSCs for 20 more sweeps. In 5-HT2R and M/Kv7 blocking experiments, the blocker dissolved in ACSF was first bath applied to the slice for at least 15 minutes before patching and recording any neurons. Baseline recordings in these experiments were obtained in the blocker alone (ketanserin or XE-991, 10 µM) before bath-applying the blocker in combination with 25CN-NBOH. sEPSCs were analyzed using Easy Electrophysiology (version 2.4.0). sEPSCs were analyzed in 30-second sweeps and the first second of each sweep was discarded as it contained the hyperpolarizing test pulse. Events were first identified with threshold-based detection with the following settings: negative peak direction, 5ms local maximum period, 30-ms decay search period, 8 pA threshold, 10ms search period, 1-ms averaged baseline, curved baseline and threshold. To eliminate false positives, events were manually inspected, and noise events were rejected. Frequency was determined for each sweep by diving the total number of events per sweep by 29 seconds, amplitude was computed by averaging all events for each sweep, half-width and decay time constant were determined from fitting an exponential to the averaged event for each sweep. The first three of twenty-two total sweeps were used as baseline. The last 8 sweeps were used to determine the mean in-drug sEPSCs.

#### In vivo drug injections and subsequent ex vivo electrophysiology experiments

DOI was administered via IP injection (2 mg/kg). After 48-72 hours, slices containing ACC were prepared and baseline electrophysiology experiments were performed as described above. The control group was a region-matched and age-matched subset of the baseline data from neurons with standard internal solution and no drugs bath-applied at baseline. Two 30-s sweeps were analyzed for sEPSC analysis (the first two baseline sweeps in the control case).

#### Statistics

Statistical tests were performed with GraphPad Prism (version 10.0.1). Data were tested to determine normal or lognormal distributions before being tested with parametric or nonparametric tests, respectively. Specific statistical tests used are provided in figure legends.

#### Data-assimilation and computational models

In order to construct a Hodgkin-Huxley model which both agrees with the observed experimental voltage traces, and incorporates explicit ion channel dynamics, we utilized the data assimilation approach pioneered by Abarbanel and coworkers (*62*, *63*). Briefly, this approach casts the parameter inference task as an optimal control problem. Suppose 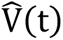 is V an experimentally observed voltage trace, and *I*(*t*) is the applied current (chosen by the experimenter). We posit a generative model given by the ordinary differential equation 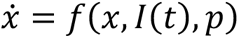 with 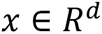, which encodes a Hodgkin-Huxley model with unknown parameters *p*. In particular, assume the first coordinate of *x* represents model voltage *V*, and the remaining *d* − 1 coordinates are model gating variables. We now desire to choose parameters *p* to minimize the discrepancy between the model voltage *V* and the observed voltage 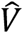; a natural optimization problem encoding this objective is 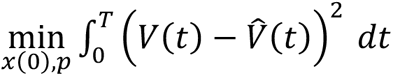 subject to the dynamic constraint 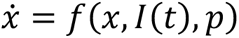. Here, [0, *T*] is the time interval over which voltage is observed, and *x*(0) is the (unknown) initial condition of the Hodgkin-Huxley dynamics. Note, however, that the above optimization problem is typically nonconvex, presenting with many spurious local minima. Instead, it is more productive to consider a regularized version of the problem, in which the model voltage dynamics incorporate a control term: 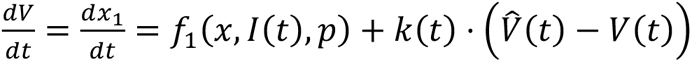 where the control *k*(*t*) ≥ 0. Let 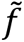 denote the controlled vector field (wherein only the first term 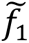, disagrees with the corresponding term *f*_1,_ in the original dynamics). Observe that as *k* → ∞, the model voltage *V* will be forced to match 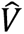; we prefer *k* → 0, so that the controlled model dynamics recover the true model dynamics.

Accordingly, we solve the optimal control problem min 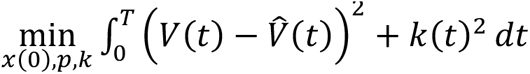 subject to the controlled dynamic constraint 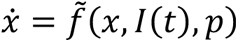. If the Hodgkin-Huxley model is appropriate to describe the data, then we hope to find solutions to this problem for which 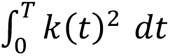 is close to zero; conversely, if no such weakly controlled solutions exist, then it may be necessary to update the generative model.

In practice, we solved this optimal control problem using a direct method (*123*), transcribing the dynamics using a fixed-order Simpson-Hermite discretization. We computed the requisite derivatives analytically using Julia code, which interfaces with the symbolic computation library SymPy. We then passed these derivatives to the interior point solver Ipopt, utilizing the sparse linear solver HSL Ma97. Heuristics for choosing the appropriate parameter bounds can be found in (*62*). We assimilated to a model incorporating the following conductances: transient sodium, delayed rectifier potassium, persistent sodium, M/Kv7 current, and H-current (parameter values in Table 6). We then pruned conductances set to zero by the optimization (in particular, the H-current is pruned, as expected for recordings coming from prefrontal IT cells (*124*) and reduced the model further by assuming infinitely fast activation kinetics for transient sodium.

In simulations including stochastic sEPSCs, sEPSC event times were generated as a homogeneous Poisson process and convolved with a first-order exponential kernel to obtain a conductance time series. The rate of the Poisson process, decay time of the exponential kernel, and maximum conductance of the simulated excitatory channel were chosen such that resultant EPSP magnitude and kinetics aligned with experimental observations. Resting membrane potential depolarization was implemented simply as application of a tonic background current having appropriate amplitude to effect the desired depolarization.

To simulate spiking-induced persistent firing, we added a model of calcium-activated nonselective cationic current (I_CAN_) to the assimilated model. I_CAN_ is represented as a slow-activating current (*τ_activation_* = 1.5 seconds) with a reversal potential of 0mV whose steady-state activation is entirely dependent on intracellular Ca^2+^ concentration, [Ca^2+^]_in_. Each time the initiation of a new action potential is detected (defined as dV/dt exceeding 10mV/ms), a unitless variable representing [Ca^2+^]_in_ is increased by 1. [Ca^2+^]_in_ slowly decays towards zero over time (τ*_decay_* = 3 seconds). To simulate elevated cholinergic tone, we reduced M/Kv7 current to 50% conductance. To simulate psychedelic effects on glutamate release, sEPSC frequency was matched to the experimentally observed data as described above.

#### Cell filling, confocal microscopy, and morphological reconstructions

To reveal cell morphology, biocytin (0.5%; Sigma Aldritch) was added to the intracellular solution. Biocytin was allowed to diffuse into the cell for no less than 15 minutes. At the end of recording, the patch pipette was retracted from the cell slowly to improve the membrane resealing, and then the slice was transferred from the recording chamber to 4% PFA for at least overnight fixation. Afterwards, each slice was washed in PBS and incubated for 24–48 hours in streptavidin conjugated Alexa Fluor (488 or 647; Thermo Fisher Scientific) with 0.4% Triton X-100 (Simga Aldritch). To visualize cytoarchitecture and aid laminar demarcation, fluorescent nissl stain, NeuroTrace (435/455; Thermo Fisher Scientific), was added to the Alexa Fluor incubation step (at 1:200 dilution). After incubation, slices were washed in PBS, mounted on slides, and cover slipped using Fluoromount-G. Slides (Southern Biotech) were left to set for 24h before being imaged. Cell imaging was done with the Zeiss Axio Image M2 microscope equipped with a LSM 700 confocal system (Zeiss). For each cell, a full zoomed out image showing the location of the cell in its entire cortical area (prelimbic or anterior cingulate) was acquired using a dry 20x objective. All such images were dual channel, one for the filled cell, and one for NeuroTrace for layer identification. These images were used for measuring distances from the border of L1/2 to the recorded neuron cell body. A subset of cells that showed strong filling with no major branches cut off were further selected as candidates for reconstruction. Z-stacks of those cell were acquired using an oil 63x objective and reconstructions were performed manually using NeuTube software (*125*).

**Fig. S1.**
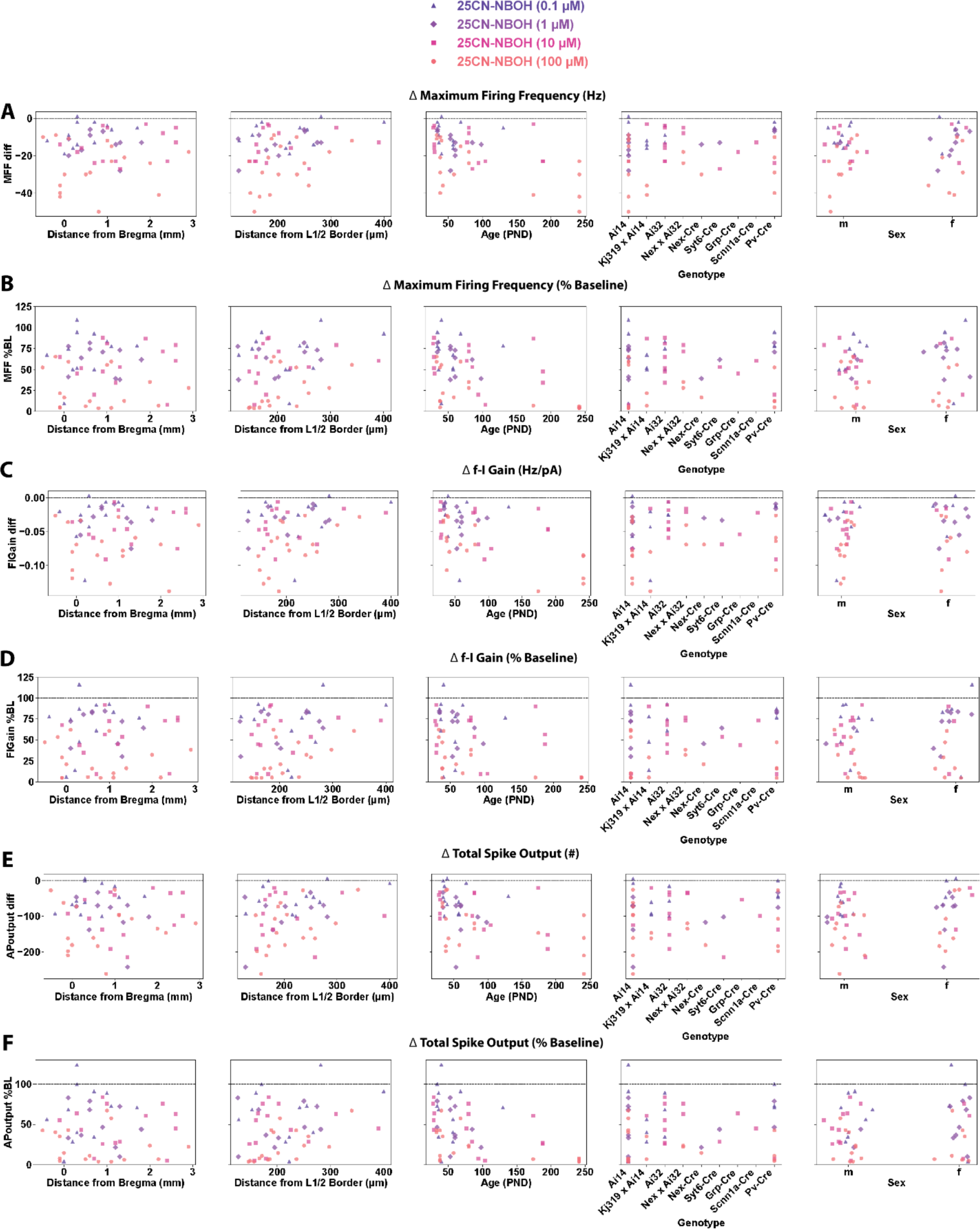
Spike suppression throughout somatic positions, ages, genotype, and sexes. Dose-dependent suppression of intrinsic excitability, as determined by evoked spiking output, was observed in PFC L5 PCs throughout: anteroposterior position within PFC (column 1), laminar depth (column 2), age (column 3), genotype (column 4), and sex (column 5). Data are shown normalized with baseline subtraction (‘diff’; A, C, E) and % of baseline (‘%BL’; B, D, F). Note that intrinsic excitability is below baseline for all doses (0 for ‘diff’; 100 for ‘%BL’), with the exception of 1 cell at the lowest dose.

**Fig. S2.**
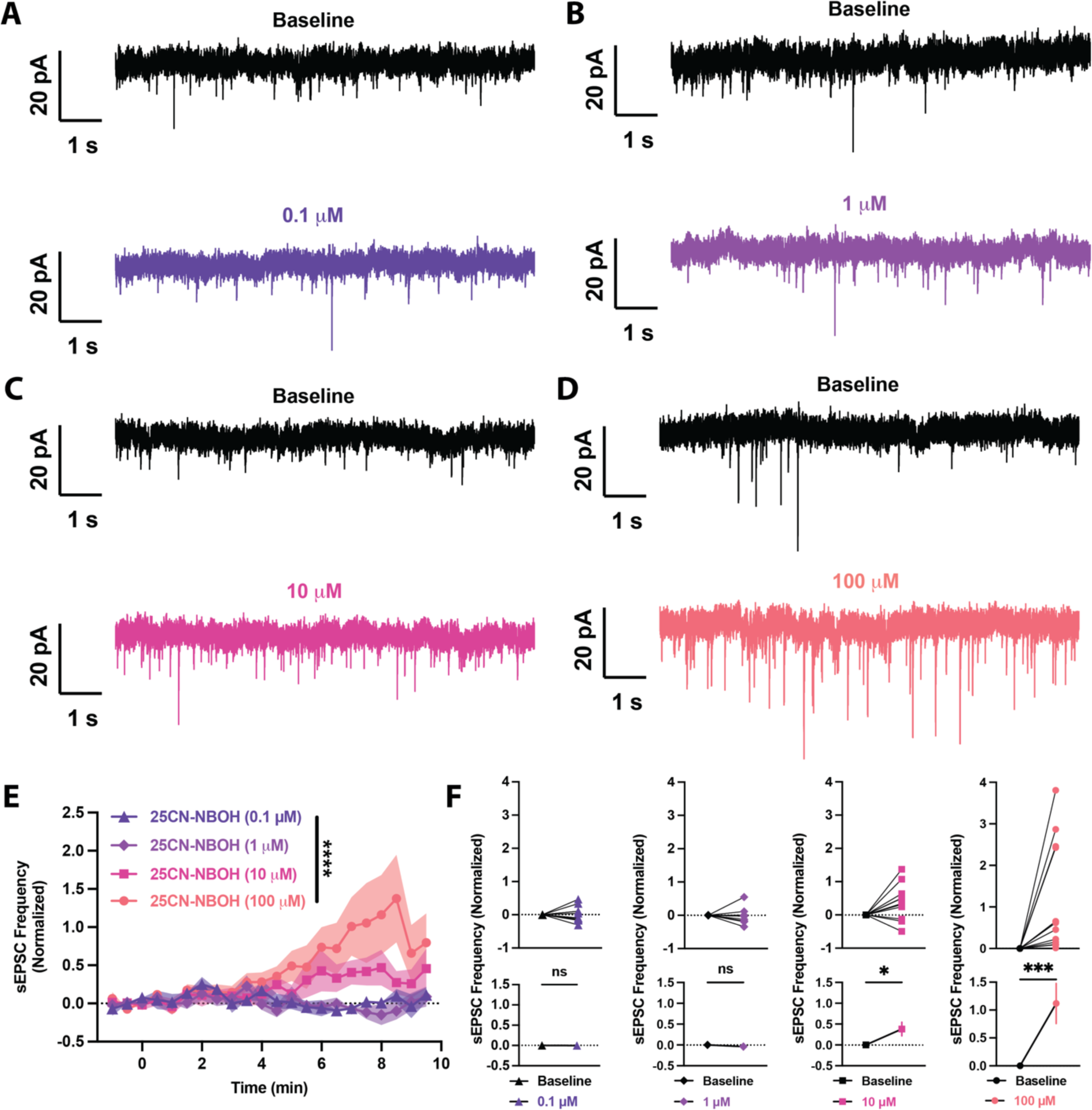
25CN-NBOH enhances spontaneous glutamate release. (A to B) Individual examples of sEPSCs during baseline (top) and in drug (bottom) for 0.1 µM (A), 1 µM (B), 10 µM (C), 100 µM (D) NBOH. (E) NBOH induces a thresholded, dose-dependent, sustained elevation of sEPSC frequency. (F) Individual cells (top) and mean data (bottom) showing that sEPSC frequency was significantly increased at 10 and 100 µM NBOH, with the strongest enhancement at 100 µM. NS, not significant; *p<0.05; **p<0.01; ***p<0.001; ****p<0.0001, F-test between simple linear regressions of dose groups (E), two-tailed paired t test between baseline and average drug response (F). Error bars and shaded regions represent (SEM).

**Fig. S3.**
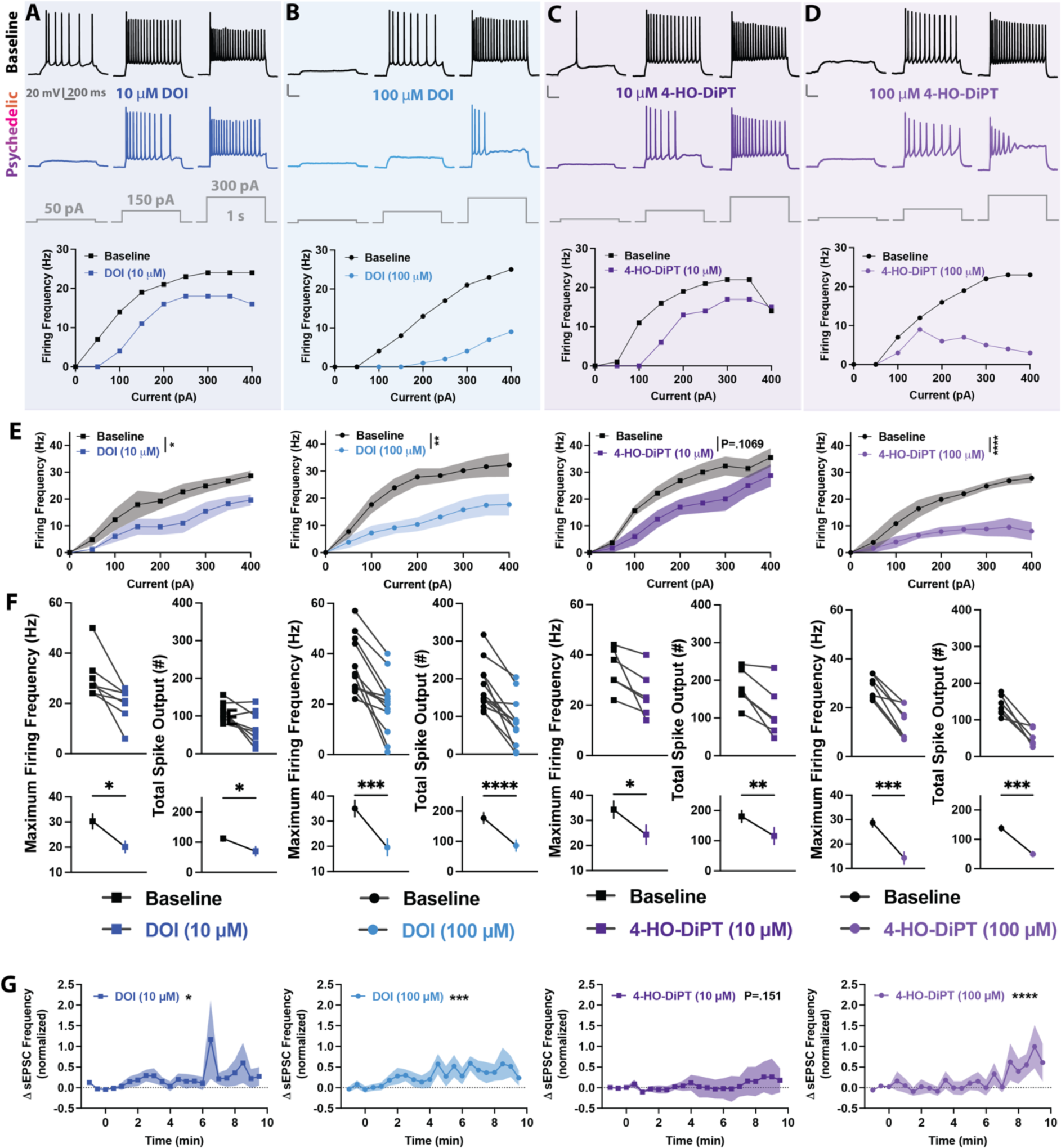
Psychedelic drugs of different classes dose-dependently suppress intrinsic excitability and enhance synaptic excitability. (A to D) Individual PFC L5 PC firing patterns (top) and f-I curves (bottom) during baseline (black) and after 10-minute bath perfusion of the psychedelic phenethylamine DOI (blue) or the psychedelic tryptamine 4-HO-DiPT (purple): 10 µM DOI (A), 100 µM DOI (B), 10 µM 4-HO-DiPT (C), 100 µM 4-HO-DiPT (D). (E) Mean f-I curves indicating dose-dependent suppression of spiking by DOI and 4-HO-DiPT. (F) Individual cells (top) and mean data (bottom) of 10 and 100 µM DOI and 4-HO-DiPT showing suppression of maximum firing frequency and reduction of total spike output with both doses of each drug, and stronger suppression with the highest doses. (G) DOI and 4-HO-DiPT dose-dependently increase frequency of sEPSCs. *p<0.05; **p<0.01; ***p<0.001; ****p<0.0001, F-test between simple linear regressions of baseline and drug (E), two-tailed paired t test between baseline and drug (F), simple linear regression.

**Fig. S4.**
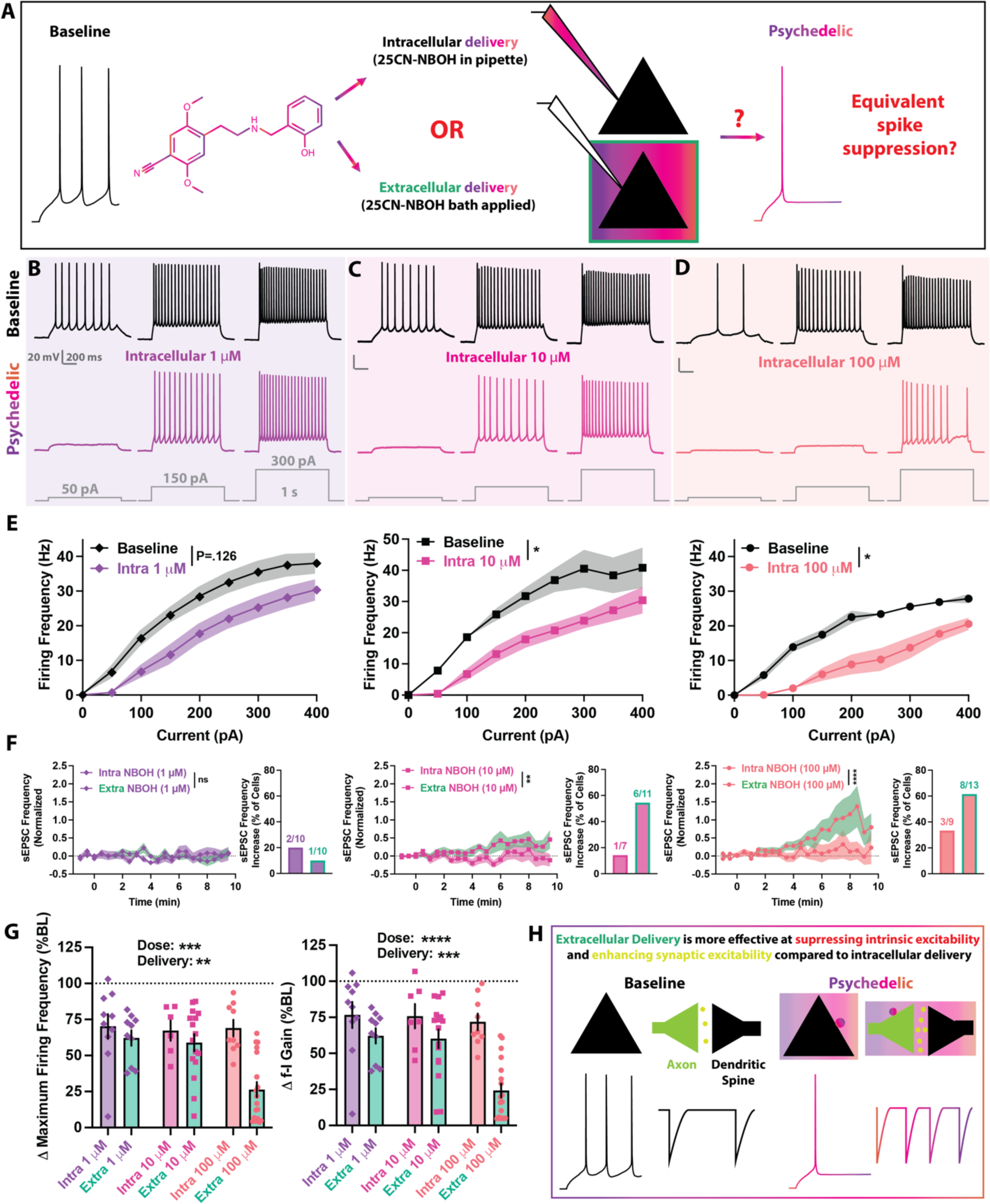
Extracellular delivery of psychedelics is significantly more effective at suppressing intrinsic excitability and enhancing synaptic excitability compared to intracellular delivery. (A) Experimental theory, design, and prediction. To strongly activate intracellular 5-HT2ARs NBOH will be added into the internal solution of the recording electrode. Firing properties after a 10-minute ‘wash-in’ period will be compared to baseline firing properties. If intracellular 5-HT2ARs mediate the radical intrinsic excitability suppression, then intracellular delivery of psychedelics will cause excitability suppression comparable to an equivalent concentration bath-applied. (B to D) Individual PFC L5 PC firing patterns during baseline (black) and after 10-minute ‘wash in’ period: 1 µM Intracellular NBOH (B), 10 µM Intracellular NBOH (C), 100 µM Intracellular NBOH (D). (E) Mean f-I curves indicating moderate suppressing of spiking in response to intracellular NBOH, but unlike extracellular delivery, lacking dramatic suppression at 100 µM. (F) Intracellular NBOH is significantly weaker at enhancing sEPSC frequency as compared to extracellular delivery. The sEPSC frequency enhancement was weaker and the percentage of cells with significantly increased sEPSC frequency was lower than with extracellular delivery. (G) Intracellular NBOH is significantly weaker at suppressing evoked spiking as compared to extracellular delivery. Changes to maximum firing frequency and f-I gain were significantly impacted by both dose, as previously shown, and delivery method. (H) Extracellular delivery of psychedelics promotes stronger changes to intrinsic and synaptic excitability than an equivalent concentration applied intracellularly. NS, not significant; *p<0.05; **p<0.01; ***p<0.001; ****p<0.0001, F-test between simple linear regressions of baseline and drug (E), F-test between simple linear regressions of intracellular and extracellular delivery (F), two-way ANOVA for dose and delivery method (G). Error bars and shaded regions represent (SEM).

**Fig. S5.**
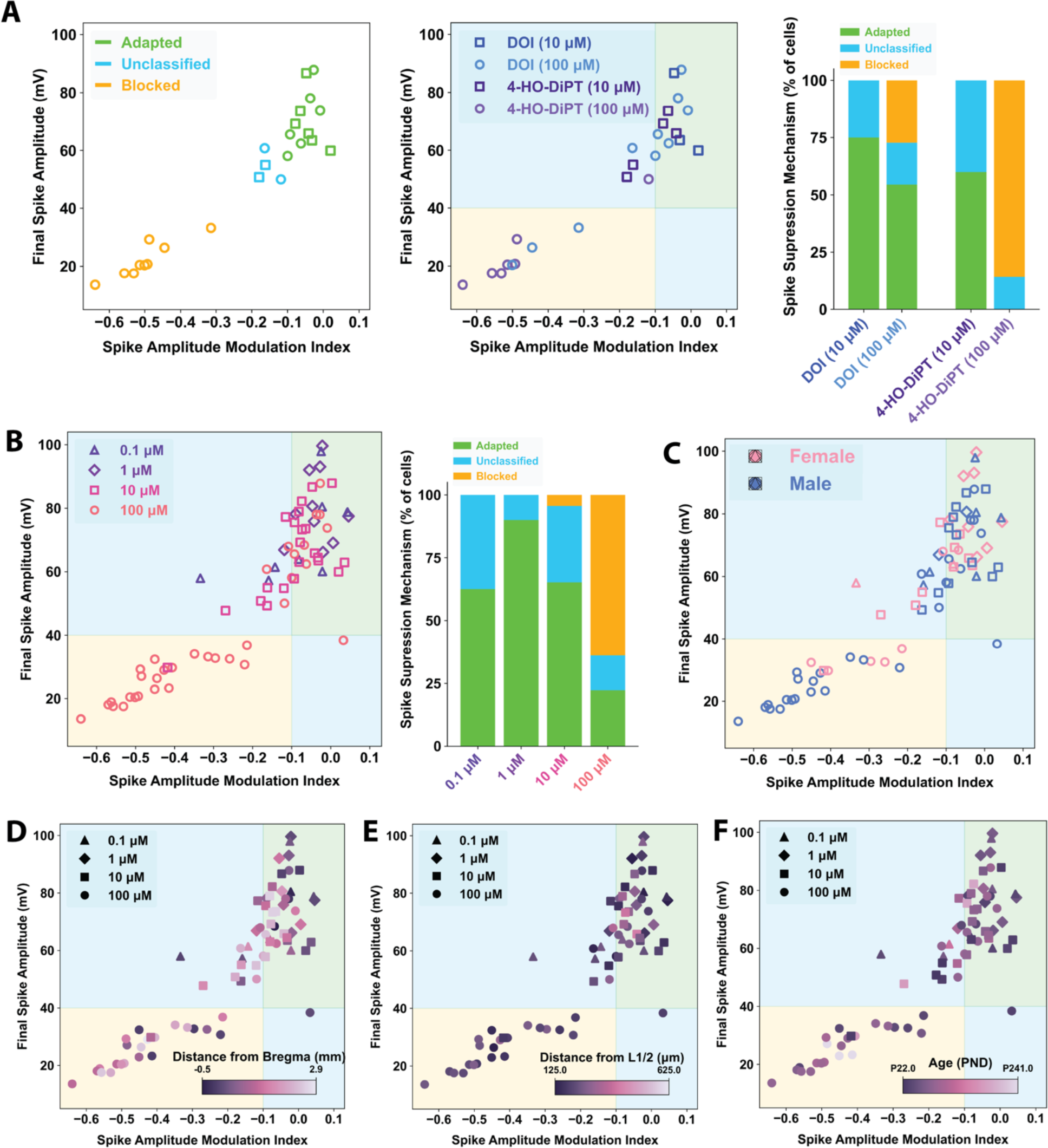
Dose-dependent spike amplitude modulation by psychedelic drugs occurs throughout sexes, somatic positions, and ages. (A) Spike amplitude modulation plots and summary plots for DOI and 4-HO-DiPT showing more adaptation at 10 µM and more depolarization block at 100 µM. (B) Spike amplitude modulation (left) and dose summary plots (right) pooling all extracellular NBOH, DOI, and 4-HO-DiPT by dose. (C to F) Spike amplitude modulation plots overlaying sex (C), anteroposterior position within PFC (D), laminar depth (E), and age (F). Symbols indicate dose.

**Fig. S6.**
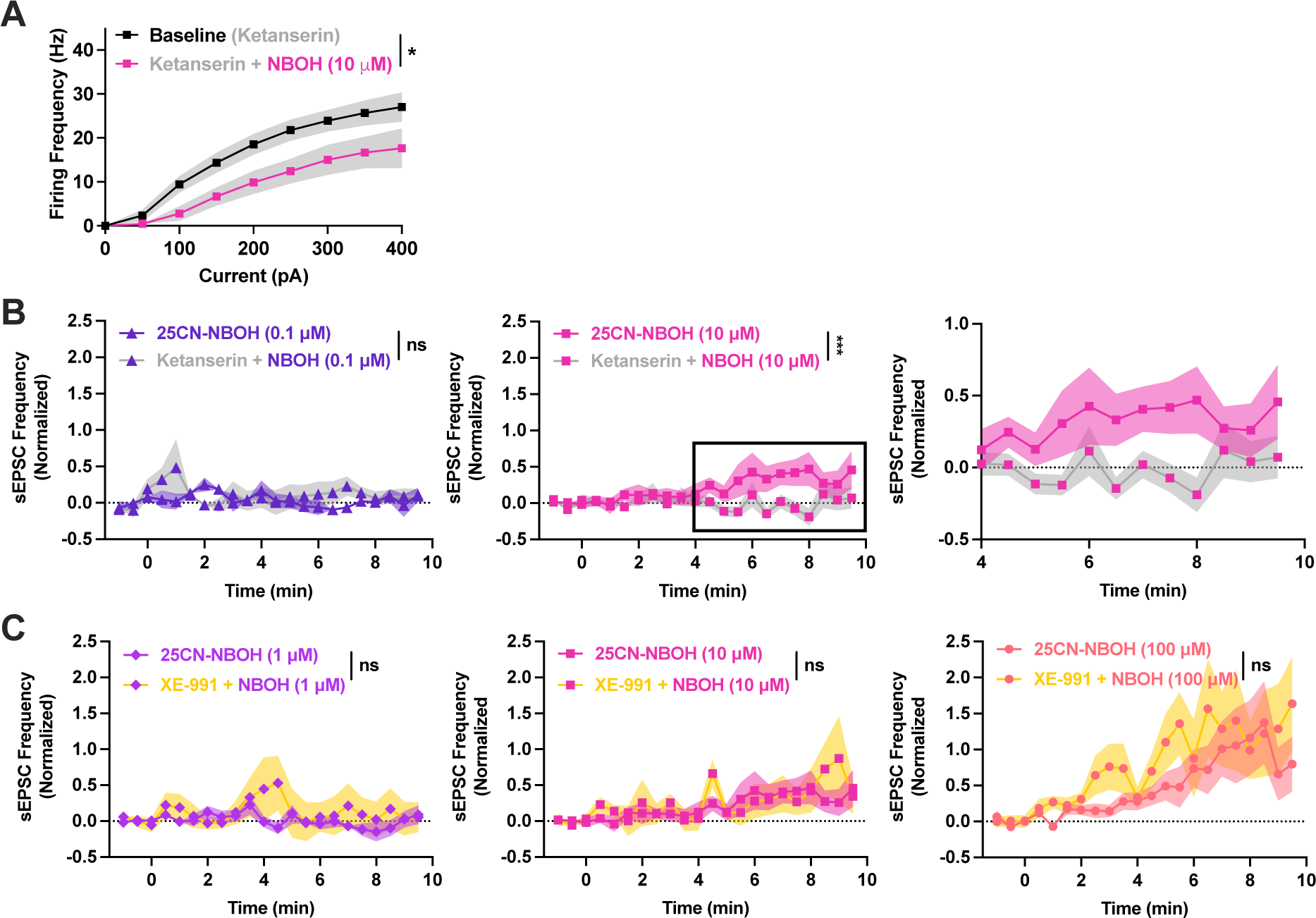
Psychedelic enhancement of sEPSCs is dependent on 5-HT2Rs but not M-current. (A) Ketanserin (10 µM) pretreatment fails to prevent fully prevent NBOH (10 µM) from suppressing evoking spiking. (B) 0.1 µM NBOH does not increase sEPSCs in presence or absence of ketanserin (left). Ketanserin (10 µM) pretreatment prevents 10 µM NBOH from increasing sEPSCs (middle, right). (C) XE-991 (M-current/Kv7 blocker; 10 µM) pretreatment does not prevent NBOH (10 and 100 µM) from increasing sEPSCs. NS, not significant; *p<0.05; **p<0.01; ***p<0.001, F-test between simple linear regressions of baseline (ketanserin) and drug (ketanserin + NBOH; A), F-test between simple linear regressions of NBOH and ketanserin (B) or NBOH and XE-991 + NBOH (C). Error bars and shaded regions represent (SEM).

**Fig. S7.**
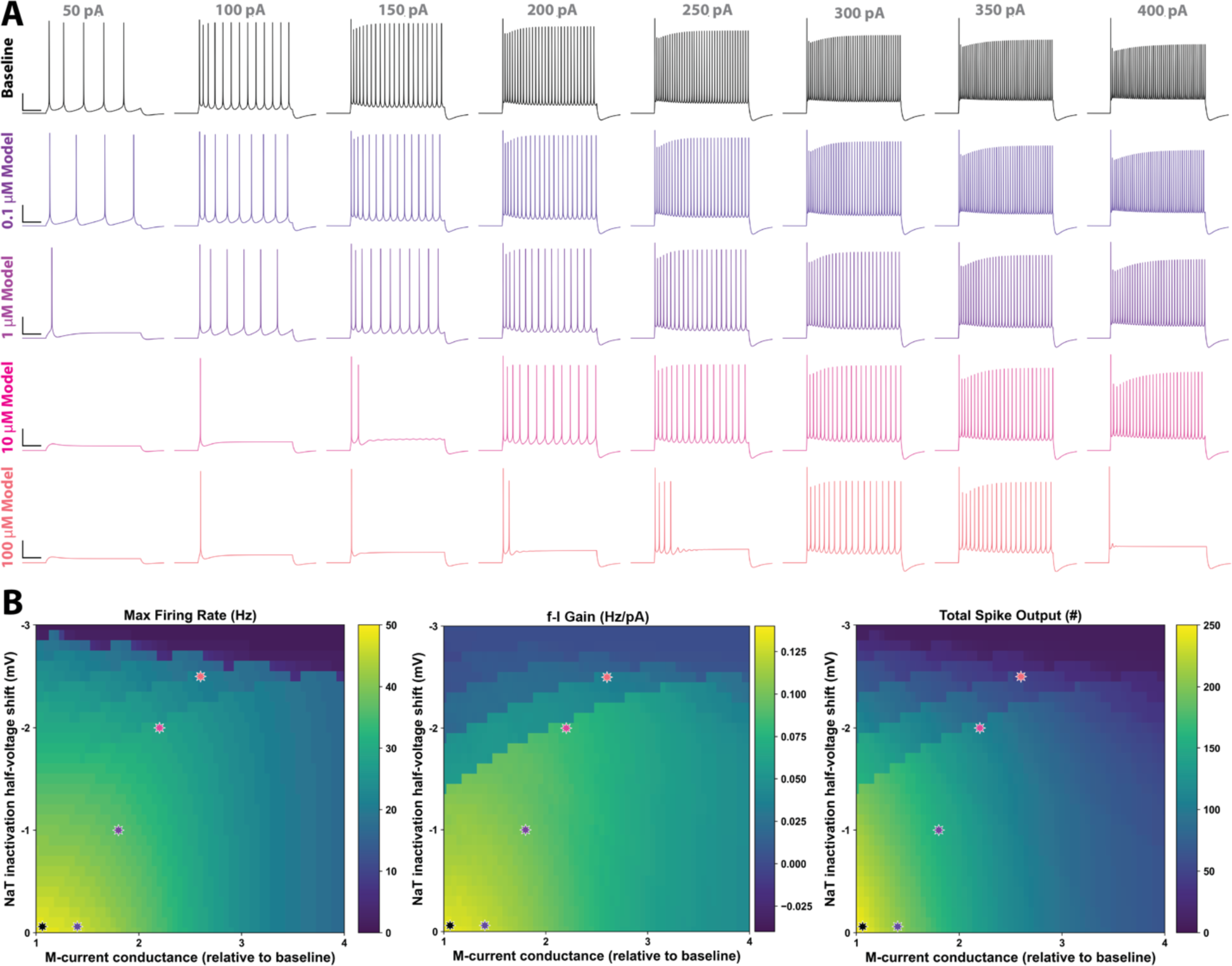
Changes to intrinsic physiology from modulations of transient sodium and M-current. (A) Firing traces of the minimal neuron model at baseline and the 4 proposed psychedelic model doses. Note occurrence of depolarization block at the 400 pA current injection of the 100 µM model. (B) Modulation of M/Kv7 and NaT in the assimilated model neuron mimics psychedelic effects on spiking output of PFC L5 PCs. Stars represent proposed model doses of 25CN-NBOH.

**Fig. S8.**
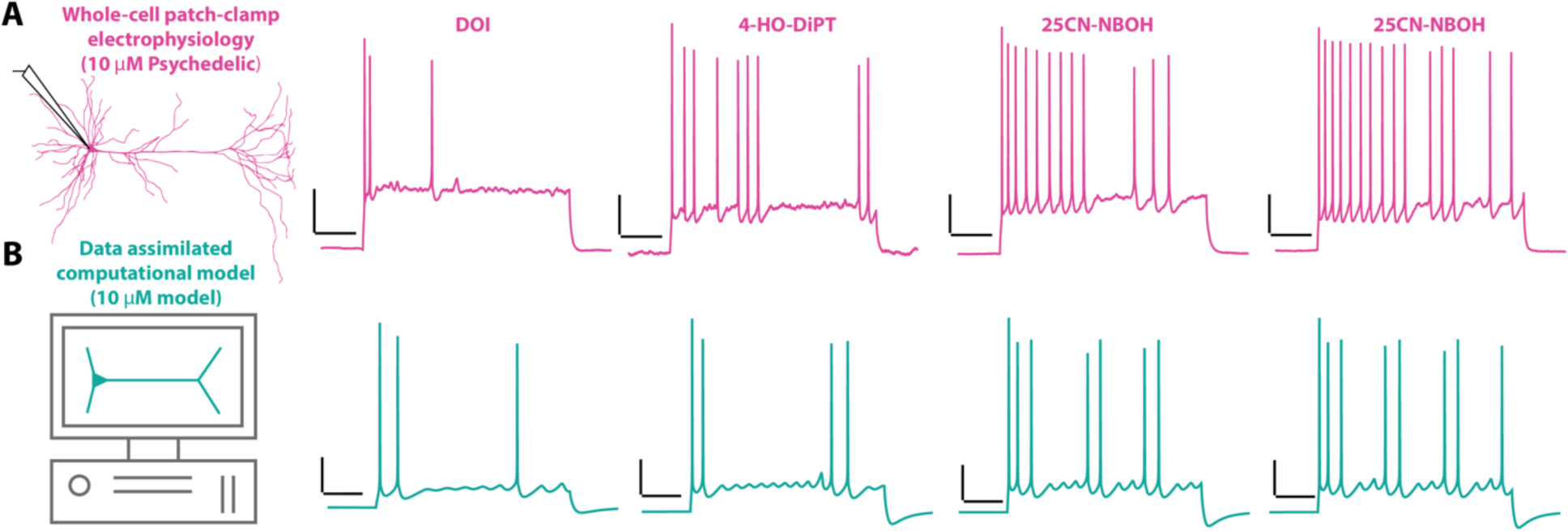
Mixed-mode oscillations from moderate levels of psychedelic activation. (A) Examples of mixed-mode oscillations in recorded neurons resulting from 10 µM application of various serotonergic psychedelics. (B) Examples of mixed-mode oscillations in the assimilated cell under M/NaT conductance modulations in and around the 10 µM dose model.

**Fig. S9.**
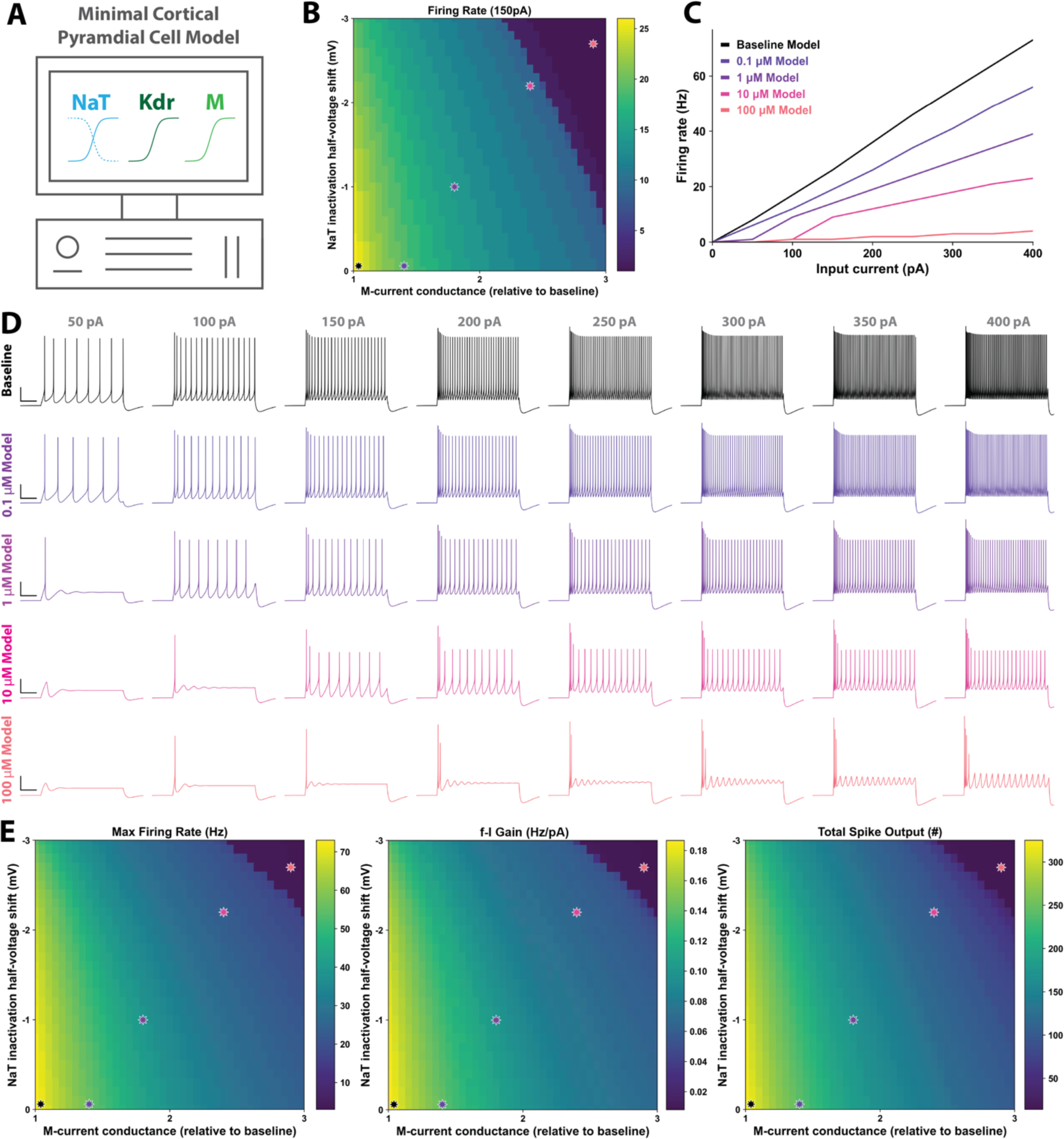
Psychedelic modulation recapitulated in a minimal cortical neuron model. (A) The Steifel model is a minimal cortical pyramidal cell model consisting only of transient sodium (NaT), delayed rectifying potassium (Kdr), and M-current (M). (B) Similar manipulations to NaT, M-current, or both reduces spiking output. Data shown for 150 pA depolarizing step pulse (1 s). (C) f-I curves of the minimal model at baseline and the 4 psychedelic dose models. (D) Firing traces of the minimal neuron model at baseline and the 4 psychedelic dose models. Note the pronounced depolarization block with the 100 µM dose model. (E) Modulation of NaT and M-current in the minimal model neuron reproduces psychedelic effects on spiking suppression. Stars represent proposed dose models of 25CN-NBOH.

**Fig. S10.**
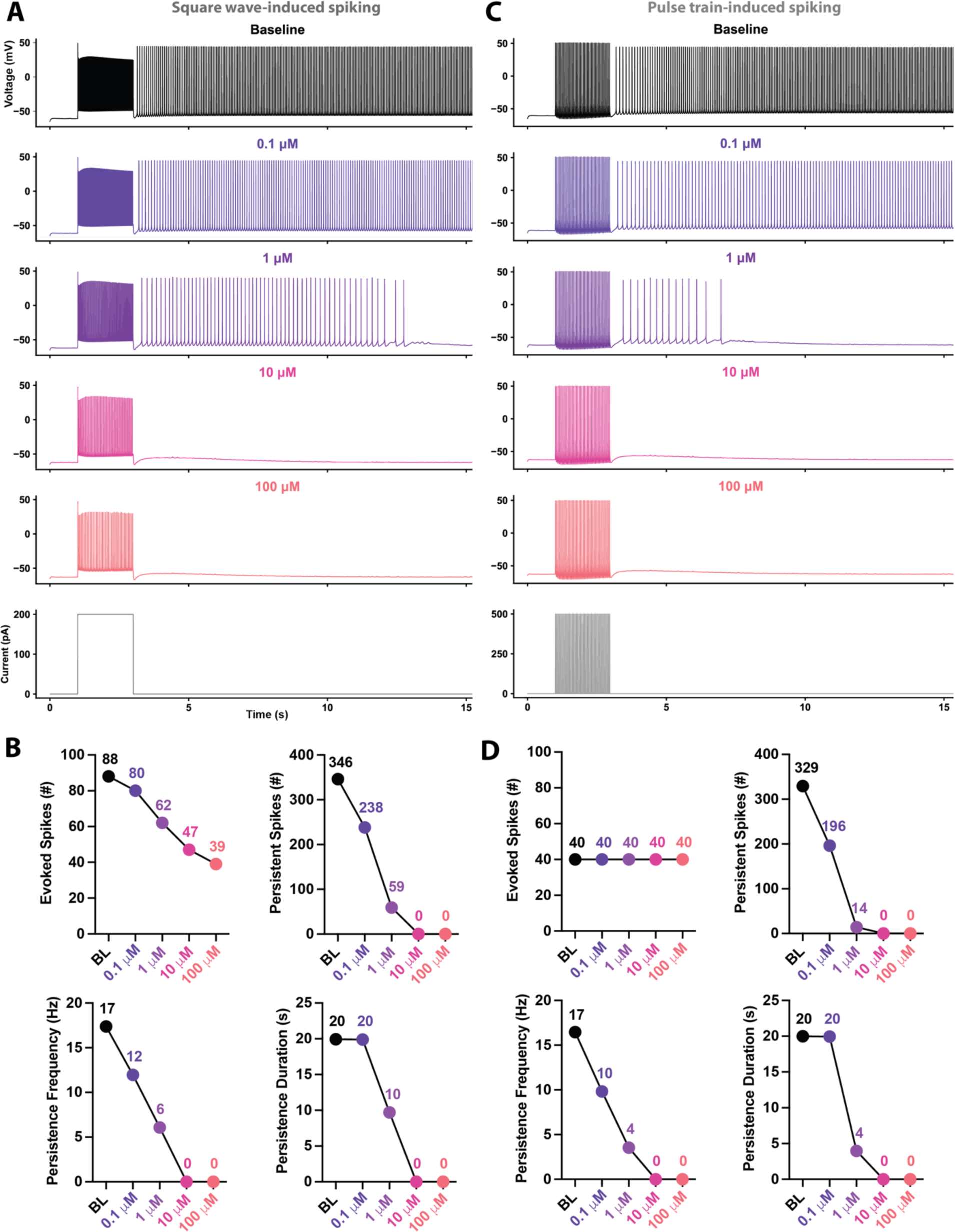
Psychedelic modulation disrupts cellular correlates of persistent working memory. (A) Square wave-induced spiking results in long-lasting ‘persistent firing’ upon addition of I_CAN_ to the assimilated model neuron. (B) Psychedelic modulation dose-dependently disrupts persistent spiking, reducing the number of persistent spikes, the firing frequency during persistence, and the duration of persistence. (C) Persistent firing occurs upon pulse train-induced spiking. 2s of 20 Hz spiking results in baseline persistence that resembles square wave-induced persistence. (D) Psychedelic modulation dose-dependently disrupts persistent spiking, even when controlling for number of spikes to induce persistence.

**Fig. S11.**
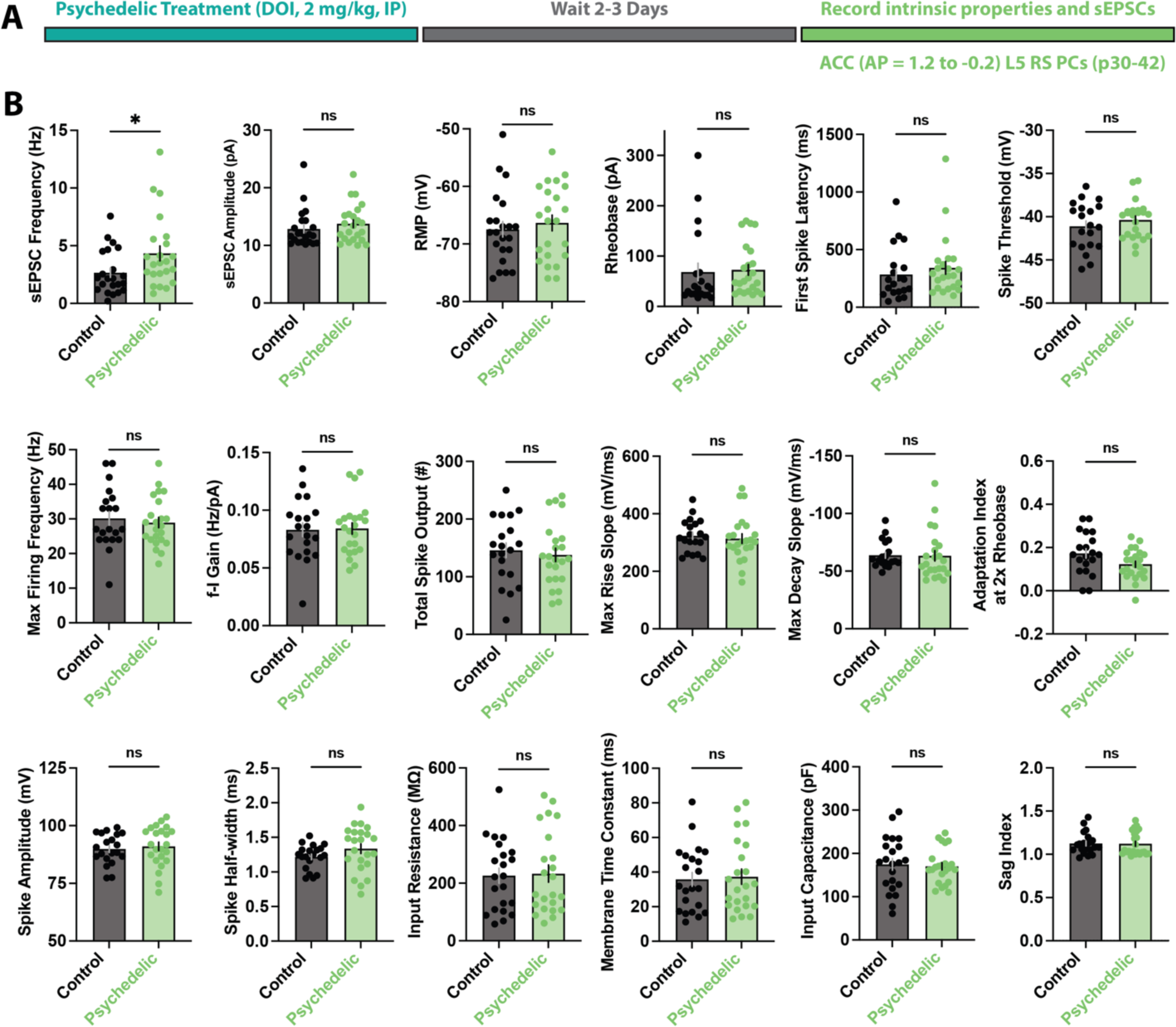
In vivo injections of psychedelics result in lasting changes to synaptic but not intrinsic excitability. (A) Experimental timeline and overview. (B) A single psychedelic treatment induces a lasting sEPSC frequency elevation but has no lasting effects on intrinsic properties. NS, not significant; *p<0.05, two-tailed Mann Whitney test between matched control and psychedelic treated groups. Error bars represent (SEM).

**Fig. S12.**
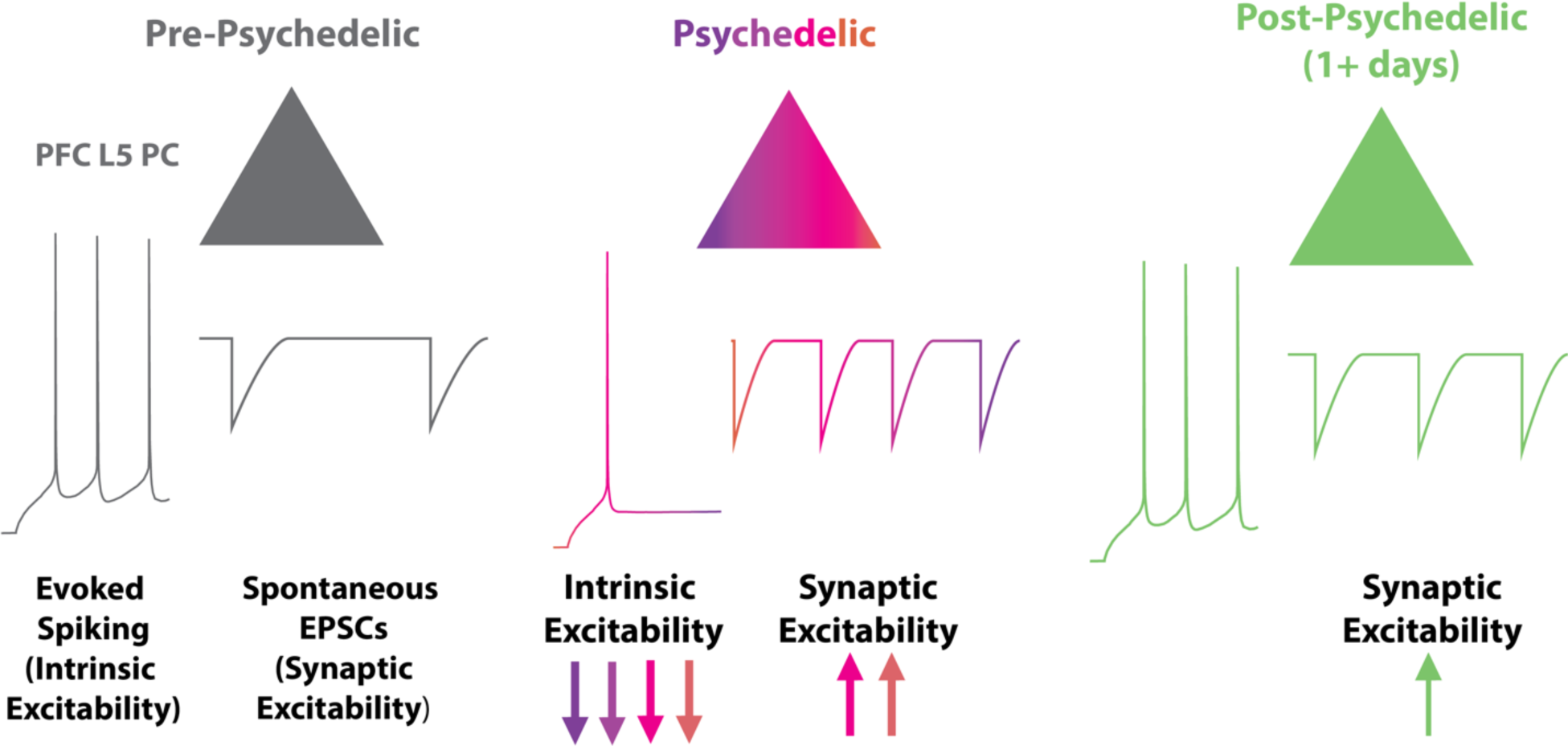
Working Model of acute and sustained psychedelic control of PFC excitability. Left, Representation of a PFC L5 PC body (top), evoking spiking (middle), and spontaneous EPSCs (bottom) in baseline conditions. Middle, the same cell in the acute psychedelic condition. Psychedelics cause a dose-dependent decrease in intrinsic excitability and a thresholded dose-dependent increase in sEPSC frequency. Right, the same cell 1-3+ days post psychedelic dose. The intrinsic excitability remains unchanged from baseline, but the synaptic excitability is higher than baseline (yet lower than the acute psychedelic state).

**Table S1.**
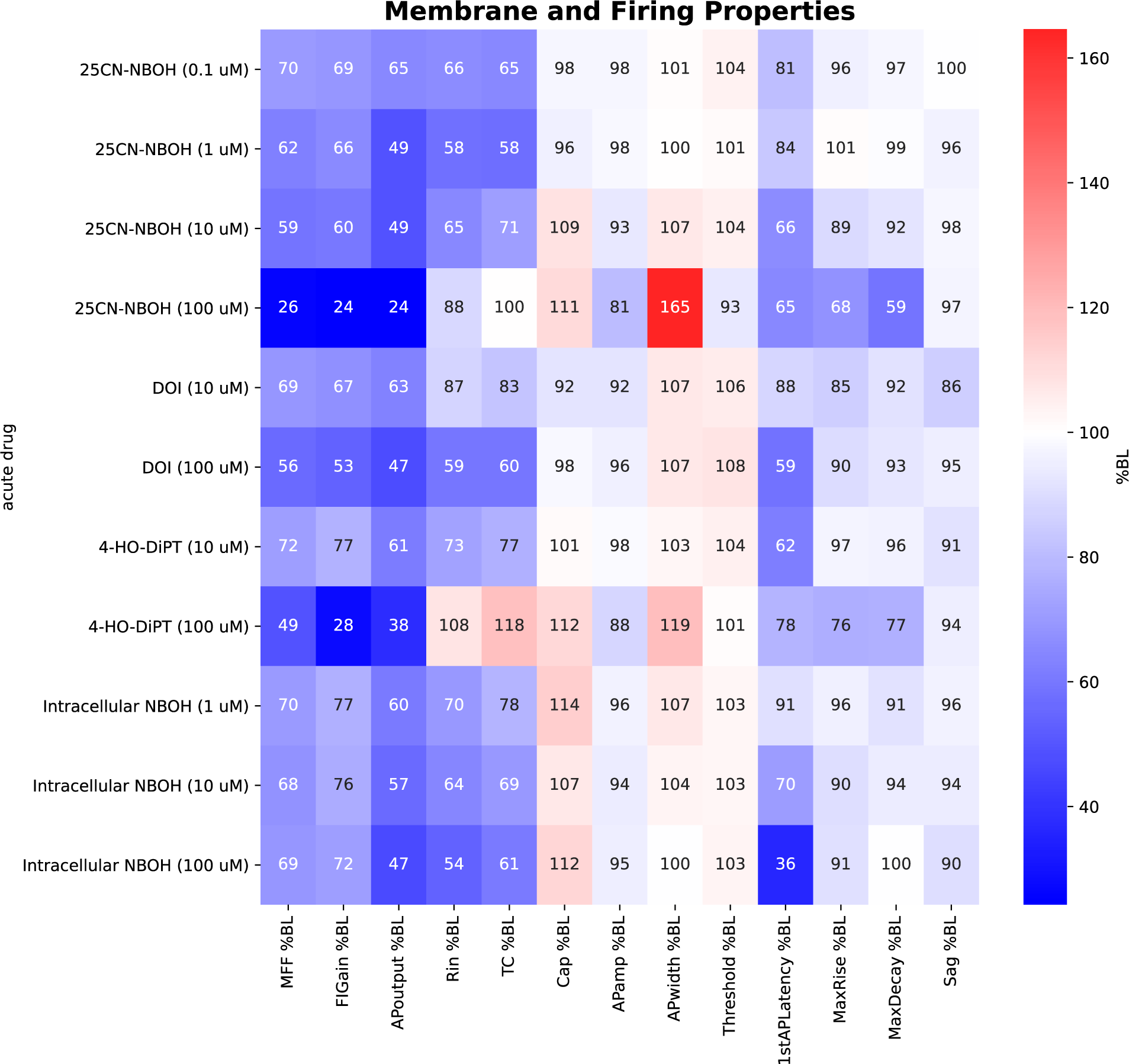
Effects of psychedelics of passive and active membrane properties. Intrinsic properties are presented as % of baseline (%BL) and are color coded to represent relative changes increased (red) or decreased (blue) relative to BL. MFF, maximum firing frequency; FIGain, frequency-current gain; APoutput, total action potential/spike output; Rin, input resistance; TC, membrane time constant; Cap, input capacitance; APamp, spike amplitude; APwidth, spike half-width; Threshold, spike threshold; 1stAPLatency, 1^st^ Spike Latency; MaxRise, maximum spike rise slope; MaxDecay, maximum spike decay slope; Sag, sag ratio.

**Table S2.**
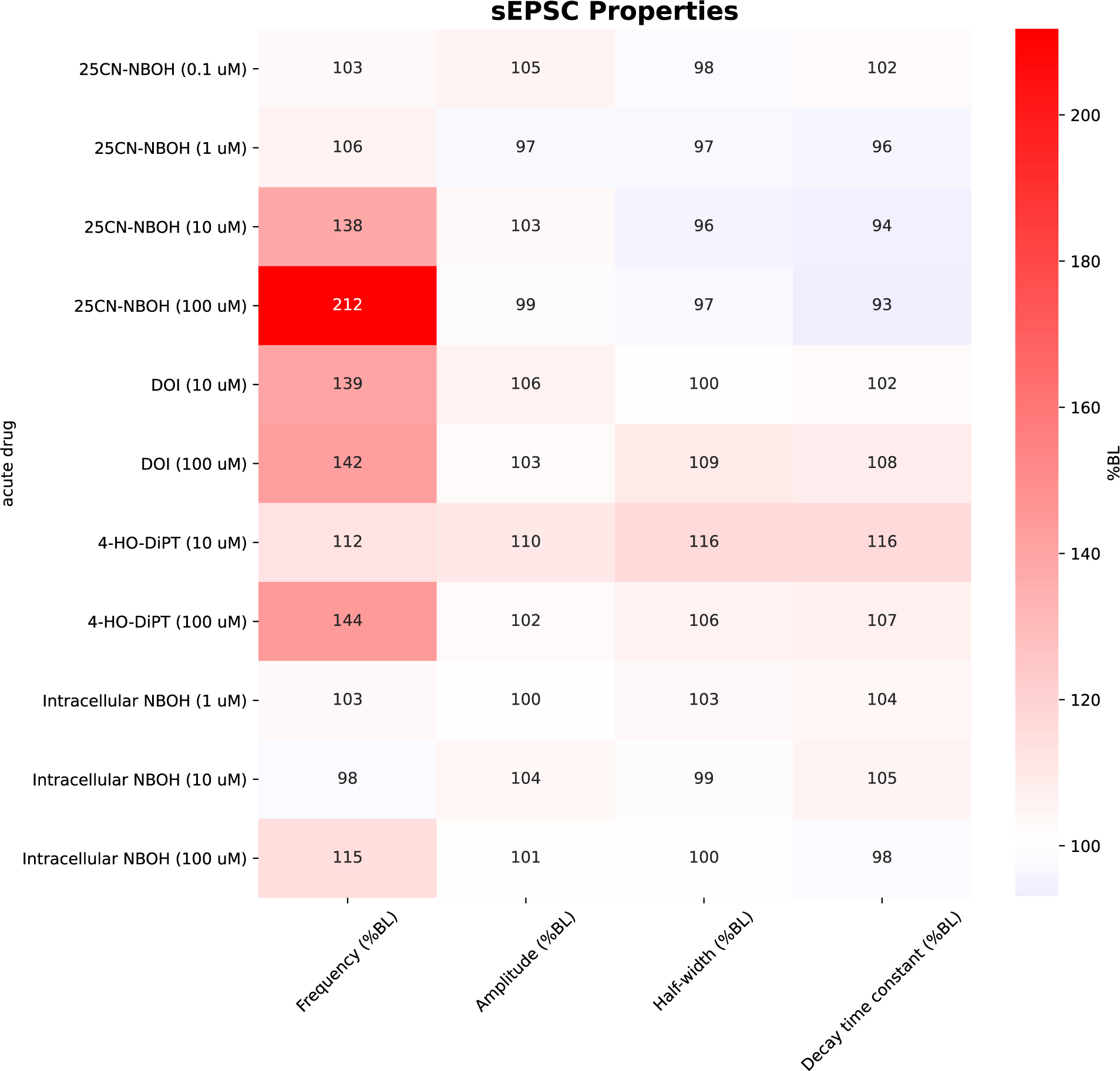
Effects of psychedelics on sEPSCs. sEPSC properties are presented as % of baseline (%BL) and are color coded to represent relative changes increased (red) or decreased (blue) relative to BL.

**Table S3.**
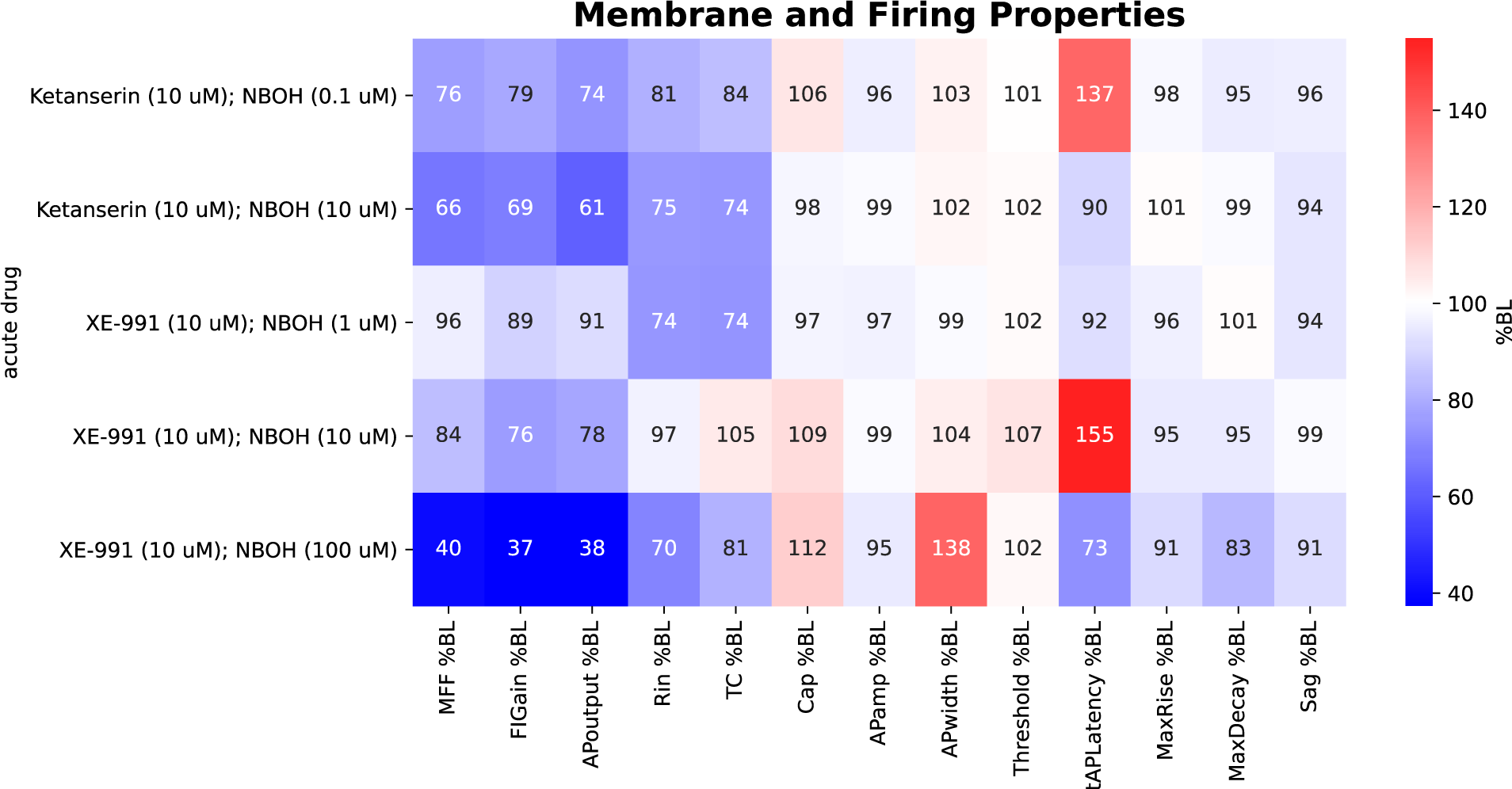
Effects of psychedelics on passive and active membrane properties in presence of ketanserin or XE-991. Intrinsic properties are presented as % of baseline (%BL) and are color coded to represent relative changes increased (red) or decreased (blue) relative to BL. MFF, maximum firing frequency; FIGain, frequency-current gain; APoutput, total action potential/spike output; Rin, input resistance; TC, membrane time constant; Cap, input capacitance; APamp, spike amplitude; APwidth, spike half-width; Threshold, spike threshold; 1stAPLatency, 1^st^ Spike Latency; MaxRise, maximum spike rise slope; MaxDecay, maximum spike decay slope; Sag, sag ratio.

**Table S4.**
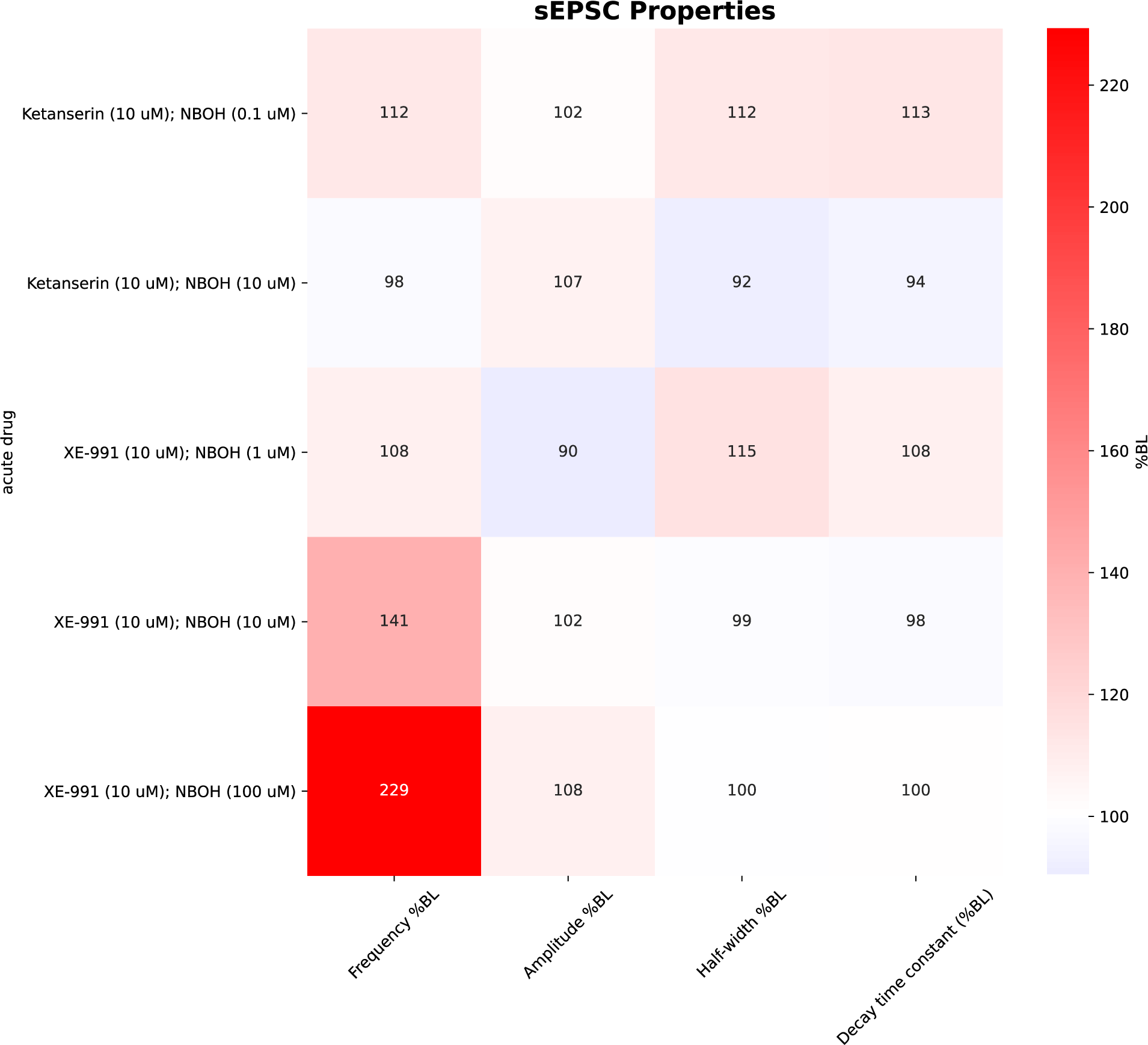
Effects of psychedelics on sEPSCs in presence of ketanserin or XE-991. sEPSC properties are presented as % of baseline (%BL) and are color coded to represent relative changes increased (red) or decreased (blue) relative to BL.

**Table S5.**
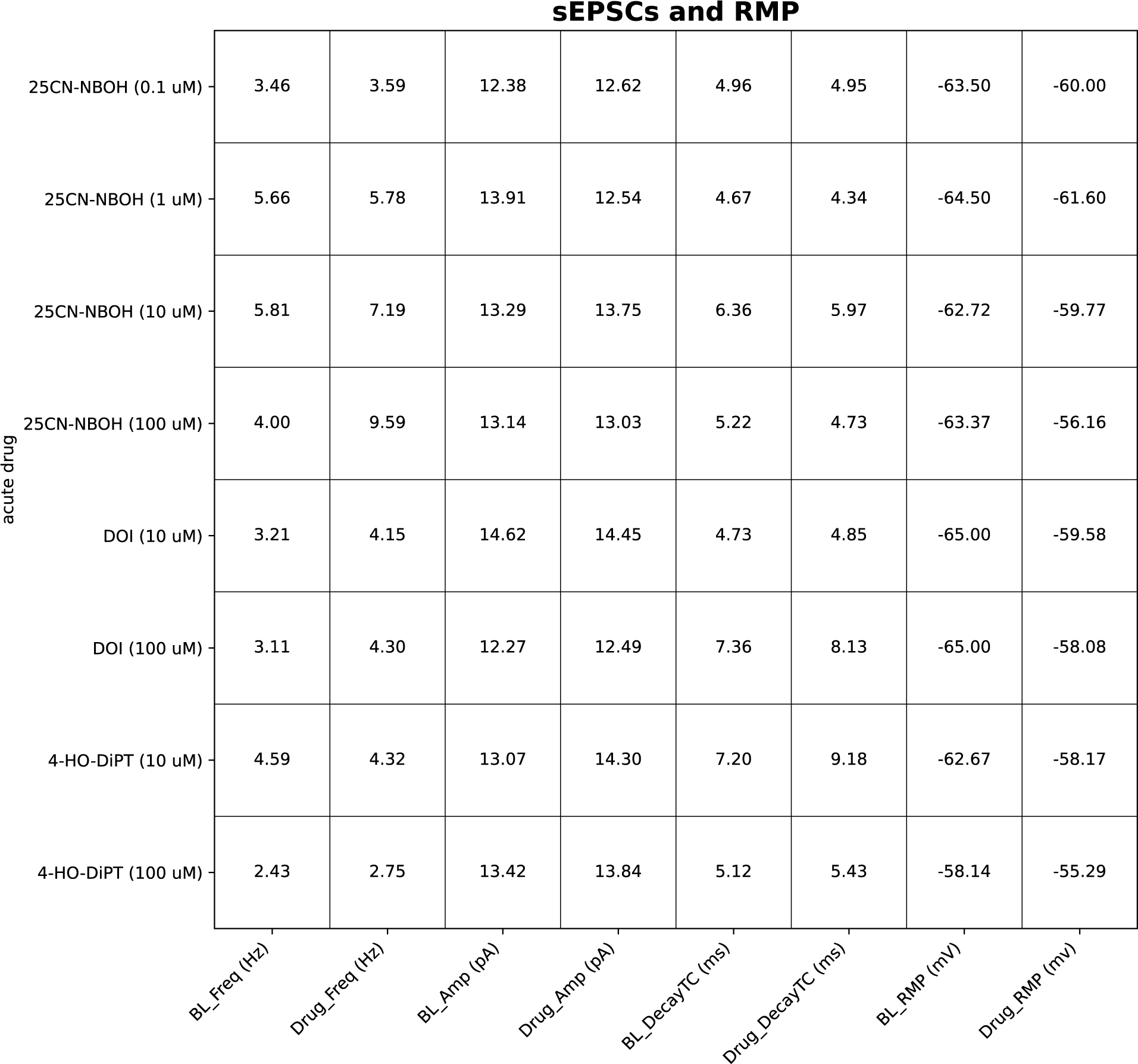
Baseline (BL) and in-drug (drug) sEPSC properties and resting membrane potential (RMP).

**Table S6.** Parameters for data-assimilated neuron.

| Variable | Value | Description |
| --- | --- | --- |
| Cm | 1.0 | Capacitance |
| I<sub>sa</sub> | 0.1 | Applied current scale factor |
| E<sub>L</sub> | -75.0 | Background leak reversal potential |
| E<sub>K</sub> | -100.0 | K<sup>+</sup> reversal potential |
| E<sub>Na</sub> | 60.0 | Na<sup>+</sup> reversal potential |
| E<sub>Glut</sub> | 0.0 | AMPA reversal potential |
| g<sub>L</sub> | 0.135 | Background leak max conductance |
| g<sub>K</sub> | 0.867 | KDR max conductance |
| g<sub>Na</sub> | 15.43 | NaT max conductance |
| g<sub>M</sub> | 0.05 | M max conductance |
| g<sub>NaP</sub> | 0.124 | NaP max conductance |
| θ<sub>m</sub> | -34.0 | NaT half-activation steady-state voltage |
| σ<sub>mss</sub> | 10.5 | NaT activation voltage-dependent slope parameter |
| θ<sub>h</sub> | -51.2 | NaT half-inactivation steady-state voltage |
| σ<sub>hss</sub> | -5.0 | NaT inactivation voltage-dependent slope parameter |
| σ<sub>htc</sub> | 15.6 | NaT inactivation time constant voltage-dependent slope parameter |
| τ<sub>h0</sub> | 0.02 | NaT inactivation time constant lower bound |
| τ<sub>h1</sub> | 12.0 | NaT inactivation time constant upper bound |
| θ<sub>n</sub> | -58.6 | KDR half-activation steady-state voltage |
| σ<sub>nss</sub> | 18.9 | KDR activation voltage-dependent slope parameter |
| σ<sub>ntc</sub> | 35.0 | KDR activation time constant voltage-dependent slope parameter |
| τ<sub>n0</sub> | 0.525 | KDR activation time constant lower bound |
| τ<sub>n1</sub> | 0.02 | KDR activation time constant upper bound |
| θ<sub>z</sub> | -38.0 | M half-activation steady-state voltage |
| σ<sub>zss</sub> | 10.0 | M activation voltage-dependent slope parameter |
| σ<sub>ztc</sub> | 11.8 | M activation time constant voltage-dependent slope parameter |
| τ<sub>z0</sub> | 0.02 | M activation time constant lower bound |
| τ<sub>z1</sub> | 400.0 | M activation time constant upper bound |
| θ<sub>p</sub> | -45.0 | NaP half-activation steady-state voltage |
| σ<sub>pss</sub> | 11.5 | NaP activation voltage-dependent slope parameter |
| σ<sub>ptc</sub> | 15.0 | NaP activation time constant voltage-dependent slope parameter |
| τ<sub>p0</sub> | 2.0 | NaP activation time constant lower bound |
| τ<sub>p1</sub> | 2.0 | NaP activation time constant upper bound |
| E<sub>ICAN</sub> | 0.0 | ICAN reversal potential |
| g<sub>ICAN</sub> | 0.015 | ICAN max conductance |
| θ<sub>ICANss</sub> | 18.0 | ICAN half-activation steady-state Ca<sup>2+</sup> concentration |
| σ<sub>ICANss</sub> | 6.0 | ICAN activation Ca<sup>2+</sup> dependent slope parameter |
| τ<sub>ICAN</sub> | 1500.0 | ICAN activation time constant |
| τ<sub>ca2</sub> | 3000.0 | Ca<sup>2+</sup> concentration decay time constant |
| spike_dca2 | 1.0 | Ca<sup>2+</sup> concentration added per spike |

